# The Ecology of Viruses in Urban Rodents with a Focus on SARS-CoV-2

**DOI:** 10.1101/2023.01.07.523115

**Authors:** Adam M. Fisher, George Airey, Yuchen Liu, Matthew Gemmell, Jordan Thomas, Eleanor G. Bentley, Mark A. Whitehead, William A. Paxton, Georgios Pollakis, Steve Paterson, Mark Viney

## Abstract

Wild animals are naturally infected with a range of viruses, some of which may be zoonotic for humans. During the human COIVD pandemic there was also the possibility of rodents acquiring SARS-CoV-2 from people, so-called reverse zoonoses. To investigate this we have sampled rats (*Rattus norvegicus*) and mice (*Apodemus sylvaticus*) from urban environments in 2020 during the human COVID-19 pandemic. We metagenomically sequenced lung and gut tissue and faeces for viruses, PCR screened for SARS-CoV-2, and serologically surveyed for anti-SARS-CoV-2 Spike antibodies. We describe the range of viruses that we found in these two rodent species. We found no molecular evidence of SARS-CoV-2 infection, though in rats we found lung antibody responses and evidence of neutralisation ability, that are consistent with rats being exposed to SARS-CoV-2 and / or exposed to other viruses that result in cross-reactive antibodies.

## Introduction

Most human infections are zoonotic and the SARS-CoV-2 / COVID-19 pandemic is the most recent example of a new zoonosis. As the size of the human population continues to increase with consequent changes in land use and greater potential for contact between animals and people then there is a continuing risk of further zoonoses (Gibb *et al*., 2020). The human population is, globally, now predominantly urban and so there is particular interest in the zoonotic potential of animals in urban environments. Comparative analysis shows that urban-adapted mammals are more parasite rich than mammal species as a whole, though not a significantly greater source of zoonoses (Albery *et al*., 2022). Rodents are already a known source of viral infections of humans. For example, hantavirsues from rodents can infect people and cause harm (Vaheri *et al*., 2013)

However, infections of people also have the potential to infect animals, so-called reverse zoonoses or zooanthroponoses. Reverse zoonoses are more likely to occur where there is substantial direct or indirect contact between people and animals and the urbanization of the human population might further facilitate this. Commensal rodent pest species, especially rats and mice, are therefore potentially susceptible to reverse zoonoses, with this more likely in urban centres where there are large populations of people and also large populations of rodents (Capizzi *et al*., 2015; Hassell *et al*., 2017). While SARS-CoV-2 is primarily a respiratory disease of people, it also infects gut tissue and viral RNA can be detected in faeces (Su *et al*., 2020; Wu *et al*., 2020) and in wastewater (Brown *et al*., 2021; Karthikeyan *et al*., 2022), and these are potential routes of transmission from people to rodents. Investigating whether commensal rodents, including those in sewers, are exposed to and / or infected with SARS-CoV-2 was the first motivation for the work that we present here.

Viruses co-infecting a host can recombine, potentially generating new viral genotypes that may have new infection characteristics, including altered host ranges (Woolhouse *et al*., 2001). A wide range of RNA viruses are already known from rats (*Rattus norvegicus*) and mice (*Apodemus sylvaticus*) (**Table 1**) and a number of rodent species are predicted to be potential hosts of SARS-CoV-2 but to also harbour other coronaviruses (*e*.*g*. Lau *et al*., 2015; Ge *et al*., 2017; Monchatre-Leroy *et al*., 2017) giving the potential for viral recombination including recombination of SARS-CoV-2 with other coronaviruses (Han *et al*., 2015; Wardeh *et al*., 2021). The second motivation for our work was therefore to discover the viruses present in commensal rodents to better understand the potential for viral recombination.

**Table 1.** A summary of the diversity and distribution of known zoonotic RNA viral families among *Rattus norvegicus* and *Apodemus sylvaticus*. Data were obtained from published articles included in the Database of Rodent-associated Viruses (DRodVir) (Chen *et al*., 2017).

| Viral family | *Rattus norvegicus* |  | *Apodemus sylvaticus* |  |
| --- | --- | --- | --- | --- |
|  | Number of viral species | Number of countries | Number of viral species | Number of countries |
| Coronaviridae | 10 | 2 | 14 | 1 |
| Arenaviridae | 12 | 2 | 39 | 2 |
| Astroviridae | 11 | 1 | 125 | 2 |
| Hantaviridae | 326 | 17 | 646 | 17 |
| Paramyxoviridae | 91 | 4 | 80 | 6 |
| Picornaviridae | 81 | 5 | 86 | 7 |

The host range of SARS-CoV-2 from early in the human pandemic excluded rats (*R. norvegicus*) and mice (*M. musculus*), though deer mice (*Peromyscus maniculatus*) and Syrian hamsters (*Mesocricetus auratus*) were susceptible (Chan *et al*., 2020; Fagre *et al*., 2021). The principal determinant of the host range of SARS-CoV-2 is the sequence of the host species’ angiotensin-converting enzyme 2 (ACE2), with which the viral Spike protein interacts to gain cell entry (Chan *et al*., 2020). During the human pandemic SARS-CoV-2 has evolved, resulting in virus genotypes with new characteristics, including altered rates of human-human transmission and an altered host range (Janik *et al*., 2021). Specifically, some SARS-CoV-2 variants of concern (VOC) are better able to infect laboratory mice compared with the original virus genotypes, showing that the host range of SARS-CoV-2 is evolving (Montagutelli *et al*., 2021). SARS-CoV-2’s host range has also been evolved experimentally: passage of human-derived SARS-CoV-2 in laboratory mice resulted in rapid evolution of viruses that better infected mice, compared with the initial human-derived virus. This change appears to be due to mutations in the viral Spike protein that newly enables cell entry through the mouse ACE2, apparently while maintaining the ability to infect through human ACE2 (Gu *et al*., 2020; Huang *et al*., 2021; Huang *et al*., 2021). One can envisage that there will be continuing selection pressure on SARS-CoV-2 to infect non-human animal species that it commonly comes into contact with, with potential effects on future human infection (Otto *et al*., 2021). In humans there are some rare ACE2 sequence variants and a similar situation may occur in wild animals, and these genotypes could be under altered selection due to sustained exposure to SARS-CoV-2 and related viruses (Shukla *et al*., 2021). The selection pressure on SARS-CoV-2 to extend its host range to rats and mice may be particularly intense in cities where there are large numbers of commensal pest rodents and also many human SARS-CoV-2 infections.

A risk assessment of the likelihood of SARS-CoV-2 infecting rodents and of onward exposure to people conducted by the UK Department for Environment, Food and Rural Affairs (DEFRA) concluded (i) with satisfactory confidence, that there was a high likelihood that a SARS-CoV-2 VOC could infect a commensal rodent, and (ii) with unsatisfactory confidence, that such rodent infections were unlikely to infect the general population but that there could be occupational exposure (DEFRA 2021). This risk assessment highlighted important knowledge gaps about SARS-CoV-2, including the (i) endogenous coronaviruses of rodents, (ii) risk of recombination of SARS-CoV-2 with other coronaviruses, (iii) selection pressure on SARS-CoV-2 to infect and transmit among rodents, and (iv) viral dose required to infect a rodent and the degree of viral shedding from any such infected animal. Recent work has sought evidence of SARS-CoV-2 in rats from sewers in Belgium, but the virus was not detected by PCR analysis, though there was some antibodies that cross-reacted with SARS-CoV-2, but these did not neutralize virus *in vitro* (Colombo *et al*., 2021). Similarly, work with *R. norvegicus* and *R. tanezumi* in Hong Kong did not find SARS-CoV-2 by PCR analysis (though this did detect other alpha and betacoronaviruses), but one rat had anti-SARS-CoV-2 antibodies and there was some evidence that these could neutralise SARS-CoV-2 (Miot *et al*., 2022). Analysis of waste water for evidence of SARS-CoV-2 has detected cryptic lineages, which are genetic variants that are not known from people, and which have been hypothesised to be SARS-CoV-2 variants infecting sewer-dwelling animals (Smyth *et al*., 2022). The Spike protein sequence of these cryptic lineages extended the *in vitro* infection phenotype to rat and mouse cells (whereas the parental lineage could only infect human cells) and was resistant to neutralization by antibodies that could neutralize the parental lineage (Smyth *et al*., 2022). The work presented here therefore further contributes to investigating commensal rodent species’ exposure and / or infection with SARS-CoV-2. In addressing this we were mindful of a number of possibilities of human-derived SARS-CoV-2 interacting with commensal rodents, ranging from full, long-lived infections, through shorter-term infections, to exposure that did not result in a patent infection.

In this work we have sampled rats (*R. norvegicus*) and mice (*A. sylvaticus*) from urban environments in 2020 during the human COVID-19 pandemic (**Figure 1**) and PCR screened them for SARS-CoV-2, metagenomically sequenced lung and gut tissue and faeces for viruses, and serologically surveyed for anti-SARS-CoV-2 Spike antibodies. We find no PCR or metagenomic evidence of SARS-CoV-2 infection, though in rats we found lung antibody responses and evidence of neutralisation ability, that are consistent with rats being exposed to and / or infected with SARS-CoV-2.

**Figure 1.**
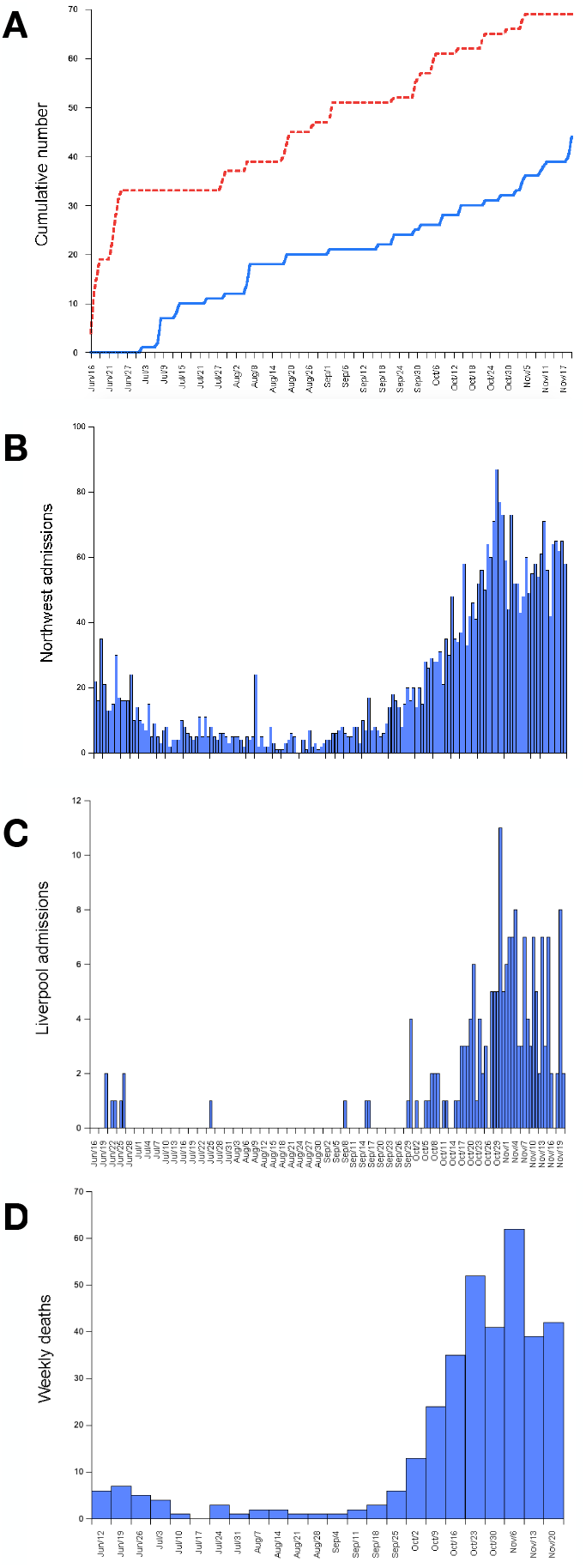
Between the 15^th^ June 2020 and 20^th^ November 2020 (A) the cumulative number of rats (blue, solid) and mice (red, dotted) caught in Liverpool, the daily number of COVID-19 hospital admissions in (B) northwest England (from https://www.england.nhs.uk/statistics/statistical-work-areas/covid-19-hospital-activity/), (C) four Liverpool hospitals (Liverpool University Hospitals NHS Foundation Trust, Liverpool Heart and Chest Hospital NHS Foundation Trust, Liverpool Women’s NHS Foundation Trust and Spire Liverpool Hospital) (from https://www.england.nhs.uk/statistics/statistical-work-areas/covid-19-hospital-activity/) (X-axis labels the same for B and C) and (D) the weekly number of COVID-19 deaths in Liverpool (from https://api.coronavirus.data.gov.uk/v2/data?areaType=utla&areaCode=E08000012&metric=newWeeklyNsoDeathsByRegDate&format=csv)

## Material and Methods

### Study species and trapping

We sampled rats (*R. norvegicus*) and mice (*A. sylvaticus*) during the human COVID-19 pandemic. The main trapping took place between 15^th^ June 2020 and 20^th^ November 2020 in and around the city of Liverpool in northwest England. We live trapped *R. norvegicus* from an urban park (Greenbank Park) and a sewage treatment works near Liverpool city centre (United Utilities, Liverpool Docks); we trapped *A. sylvaticus* from urban parks in Liverpool (Sefton Park, Princes Park, Calderstones Park and Wavertree Botanic Gardens). All animals were humanely killed with an overdose of pentobarbitone anaesthetic, and then stored at -20&C. These animals were used for viral metagenomic sequence analysis, PCR diagnosis of SARS-CoV-2, and serological analysis. An additional 41 rats (21 male, 17 female, 3 not determined) from southern England were donated from pest controllers and were sampled from a residential area in Reading (51.466479, -0.966974) and a household waste disposal facility in Basingstoke (51.278179, -1.065996); animals donated from these sources were used for serological analysis only.

### Dissection and RNA extraction

We dissected animals individually and collected tissue from the lung (all lobes) and from the small intestine, and recovered faecal material from the rectum. RNA was then separately extracted from 20 – 30 mg of these samples using the Qiagen RNeasy Mini Kit Plus following the manufacturer’s protocol and RNA samples were stored at -80&C.

### Pooling RNA samples

For metagenomic sequencing and PCR analysis we pooled RNA samples in the following way: (1) For the 10 rats caught in urban parks we pooled RNA from the same tissue type (lung, gut, or faeces) from three or four rats, with equal amounts of RNA from each rat. Pooling in this way would tell us about the tissue sample-associations of any viruses; (2) For the 35 rats caught in the sewage treatment works we pooled the three different samples (lung, gut, faeces) of each animal, with equal amounts of RNA from each sample. Pooling in this way would tell us about the prevalence of any viruses in the rat population. (3) For the 69 mice, the RNA was pooled as 2 (above) and for 35 mice this pooled RNA was sequenced for individual mice; for 34 mice RNA from randomly selected pairs of mice caught on the same day was pooled.

### Positive controls – metagenomic sequencing and PCR

For the PCR and metagenomic sequence analysis we generated positive controls by spiking tissue samples with inactivated SARS-CoV-2. Specifically, we took samples of tissue from gut, lung and faeces (as above) from laboratory SPF rats and mice (*Mus musculus*). To each of these individual samples we then generated two controls: a ‘high’ control of 10^8^ and a ‘low’ control of 10^4^ UV inactivated SARS-CoV-2 virus particles (grown in Vero E62 cells) per gram of tissue. The 10^4^ dose is one that was used to successfully infect *Peromyscus maniculatus* with SARS-CoV-2 (Fagre *et al*., 2021). These samples were then processed for RNA extraction (as above) and the RNA samples then pooled as for the rats caught in the sewage treatment works, separately for the SPF rats and mice.

### Metagenomic sequencing and bioinformatics

The RNA was rRNA depleted using the NEBNext rRNA process using human / mouse / rat baits used with the rat-derived samples, and human / mouse / rat and bacterial baits with the mouse-derived samples, following the manufacturer’s protocol. Depleted material was then purified using Ampure RNA XP beads and successful rRNA depletion was confirmed using a fragment length analyser. Depleted RNA was then used in the NEBNext Ultra II Directional RNA Library Kit for Illumina, with a fragmentation time of 7 minutes, and after 12 cycles of amplification the libraries were purified using Ampure XP beads. Each library was quantified using Qubit and the size distribution assessed. Final libraries were pooled in equimolar amounts and the quantity and quality of the pool was assessed with a Bioanalyzer and subsequently by qPCR using the Illumina Library Quantification Kit from Kapa on a Roche Light Cycler LC480II. Template DNA was diluted to 300 pM and denatured for 8 minutes at room temperature using freshly diluted 0.2 N sodium hydroxide, and the reaction terminated by the addition of 400 mM TrisCl pH 8. The libraries were sequenced on the Illumina NovaSeq 6000 platform following a standard workflow over 8 lanes of an S4 flow cell generating 2 × 150 bp paired-end reads.

Initial processing and quality assessment of the sequence data was performed using an in-house pipeline. Briefly, base-calling and de-multiplexing of indexed reads was performed by CASAVA version 1.8.2 (Illumina) to produce sample sequence files in FASTQ format that were then trimmed to remove Illumina adapter sequences using Cutadapt version 1.2.1 (Martin, 2011). The reads were further trimmed to remove low quality bases, using Sickle version 1.200 with a minimum window quality score of 20. After trimming, reads shorter than 20 bp were removed and if both reads from a pair passed this filter, each was included in the R1 (forward reads) or R2 (reverse reads) file; if only one of a read pair passed this filter, it was included in the R0 (unpaired reads) file.

We next removed host (*R. norvegicus, A. sylvaticus, M. musculus*) sequence reads from the adapter and quality-trimmed paired end read data where reference genomes were indexed and aligned with Bowtie2 (Langmead & Salzberg, 2012) and unmapped reads (*i*.*e*. reads that did not align to the host reference) were extracted. We then classified remaining reads using Kraken2 (Wood *et al*., 2019), using both the standard and viral Kraken2 database. A custom script was used to combine the Kraken2 reports into taxonomy abundance tables at different taxonomic levels.

We sought to assemble some viral genomes, which we did by taking the relevant Kraken2-defined reads and attempted genome assembly using SPAdes with default parameters (Bankevich *et al*., 2012).

For reads indicated by Kraken2 to be putatively derived from SARS-CoV-2, we mapped them using Bowtie2 and visualised their position in the genome using IGV (Thorvaldsdóttir *et al*., 2016) and compared with primer locations for ARTIC primer scheme (github.com/artic-network/primer-schemes), which was commonly used in the laboratory at the time and represents a potential source of contamination.

### Quantitative analysis

We first determined which viruses were present in each sample that was sequenced, which we did by expressing the number of Kraken2-identified reads for each virus as a proportion of all the sequence reads in that sample. This was done to control for variation in depth of sequencing among different samples. We used the results from the low positive control (10^4^ virus particles per gram of tissue) to set our detection cut-off, which was the number of SARS-CoV-2 reads as a proportion of all reads in the relevant library. We used this proportion as the lower-level cut off for other viral reads.

For each virus, we then compared this proportion to that for the low positive control, and assumed that those that were above this were positive for the respective virus. We did this separately for the rat and mouse samples and all the results presented here are after applying this threshold.

We then used these data to measure the virus infections in various way, separately for rats and mice; specifically, (i) the number of viruses, (ii) the prevalence of different viruses, (iii) the number of infections caused by different viruses, (iv) the viral load of different viruses, and (v) for rats, the tissue association of different viruses. For (i) we counted the total number of viruses in each sample, and then compared these values between rats and mice using a generalised linear model with Poisson error correction, in which host species was the fixed effect and virus number was the response variable. For (ii) we counted the number of animals that were infected by a virus and expressed this as a proportion of all the relevant animals. For (iii) for each virus (which was assigned to a viral family), we counted the number of animals that it infected, and from this determined the total number of infections caused by each viral family, which when then expressed as a proportion of the total number of viral infections. For (iv) for each virus we determined the maximum number of sequence reads for that virus for any single host, and expressed this as a proportion of all the reads that we obtained for that host. For (v) we focussed on the tissue-specific (*i*.*e*. lung, gut, faeces) pools from 10 rats, and determined which viruses were uniquely present in each tissue type.

### PCR for SARS-CoV-2

Reverse transcription and PCR screening for SARS-CoV-2 was performed on all of the pooled RNA samples. cDNA was synthesised using a LunaScript RT SuperMix Kit (New England Biolabs) and PCR was carried out using a Q5 High-Fidelity PCR Kit (New England Biolabs) following the manufacturer’s protocol. SARS-CoV-2 was amplified using the ARTIC V3 multiplex primer panel (artic.network; Artic 2020), which produces amplicons of approximately 400 bp. All reactions were analysed by gel electrophoresis.

### Serology

We used ELISAs to assay for antibodies to the SARS-CoV-2 Spike protein in wild rats and mice. We assayed tissue fluid extracts of heart, liver and lung tissue for ELISA. To do this, 0.5 g of each tissue from the wild caught rats and mice was homogenised in 1 mL of phosphate buffered saline, supplemented with Triton to a final concentration of 1 % w/v and a protease inhibitor cocktail (Sigma) to a final concentration of 20% v/v, left on ice for 30 minutes, centrifuged at 13,000 *g* for 10 minutes at 4&C, and then the supernatant removed, which was then stored at -20&C.

We generated positive control samples by immunising laboratory rats or mice (*M. musculus*) with SARS-CoV-2 Spike protein, which was based on the approach of Yang *et al*., 2020. To do this Wister rats and C57BL/6J mice were immunised intramuscularly with 5 μg SARS-CoV-2 Spike protein (residues 319-541) (Thermo Fisher) administered in TiterMax Gold Adjuvant on day 0 and on day 50. Animals were killed on day 58 and serum prepared and stored at -20&C, and tissue dissected and tissue fluid samples prepared as described above. We used non-immunised laboratory SPF rats and mice (*M. musculus*) as negative controls.

We validated the use of tissue fluid samples for ELISA in three ways. Firstly we determined the concentration of total IgG in (i) negative control laboratory rats, comparing serum, liver and heart tissue fluid samples, and (ii) a sub-sample of wild rat liver and heart tissue fluid samples. The total IgG assays were conducted for rats and mice using the IgG Total Rat Uncoated ELISA Kit (Thermo Fisher), following the manufacturer’s protocol. Secondly, we determined the concentration of anti-SARS-CoV-2 Spike IgG in positive control Spike protein immunised rats, comparing serum, liver and heart tissue fluid samples, using the relevant ELISA protocol described below. Thirdly, we determined the concentration of total IgA in (i) positive control Spike protein immunised rats, comparing serum, lung, liver, and heart tissue fluid, and (ii) a sub-sample of wild rat lung tissue fluid samples.

We used heart tissue fluid samples for IgG ELISAs and lung tissue fluid samples for IgA ELISAs. For the IgG anti-SARS-CoV-2 Spike protein ELISA, plates were coated with SARS-CoV-2 Spike protein (Thermo Fisher) at a concentration of 1 μg / mL overnight at 4&C. All buffers used were supplied by Thermo Fisher and used as recommended. After coating, plates were washed twice in wash buffer, blocked with blocking buffer for 2 hours at room temperature, then washed twice with wash buffer. Samples were diluted in assay buffer and titrated in doubling dilutions on plates, then left for 2 hours with shaking at room temperature, after which they were washed four times in wash buffer, after which a 1 : 5,000 dilution (in assay buffer) of goat anti-rat IgG horseradish peroxidase conjugate (Thermo Fisher) was added, which was incubated for 1 hour with shaking at room temperature, then washed four times with wash buffer before the addition on 100 μL of TMB substrate solution, which was then stopped after 25 minutes with stop solution. The mouse ELISAs were done in the same fashion except that goat anti-mouse IgG horseradish peroxidase conjugate (Thermo Fisher) diluted 1 : 10,000 in assay buffer was used, and that once the substrate was added the assay was stopped after 20 minutes.

We used rat lung tissue fluid samples for IgA anti-SARS-CoV-2 Spike protein ELISAs. These were conducted as described for the IgG ELISAs except that the detection antibody was goat anti-rat IgA horseradish peroxidase conjugate (Invitrogen) diluted in assay buffer at 1 : 10,000 and the reaction developed for 15 minutes before being stopped.

Note that the target mouse species is *A. sylvaticus* but the murine kits and regents are designed for use with the mouse *M. musculus*. Previous work (*e*.*g*. Jackson *et al*., 2009; Clerc *et al*., 2019) has shown cross-reactivity of antibodies between these two mouse species.

We report the ELISA results as Optical Densities (OD) or as titres. When presenting ODs we do so for a dilution of the sample where that dilution placed the ELISA data in the non-asymptotic part of the dilution *vs*. OD relationship. Titre is the reciprocal of the dilution of the serum or tissue fluid sample that achieves an OD greater than the negative in the assay, where we defined negative as the mean + 2 x standard deviations of the OD of our negative control sample.

### Neutralization assays

We tested the extent to which wild rat heart and rat lung tissue fluid could inhibit the ability of SARS-CoV-2 to infect mammalian cells, which we did using an *in vitro* neutralization assay after Adaken *et al*., 2021. Briefly, SARS-CoV-2 enveloped pseudo-typed virus particles (PVP) displaying the ancestral, Wuhan Spike protein (Accession MN908947) were generated by transfecting HEK293T Lentix cells with a pCSFLW lentiviral luciferase reporter, the SARS-CoV-2 envelope expression plasmid, and the lentiviral backbone p8.91 (Carnell *et al*., 2017; Di Genova *et al*., 2021). Samples were serially diluted, added to PVP and held at room temperature for 30 minutes. Next, this virus / sample dilution mix was used to infect HEK293T ACE2 TMPRSS2 cells and PVP infection was monitored by luciferase activity, where (i) 0 % inhibition was taken as the infection values of the virus in the absence of human convalescent plasma included in each experiment, and (ii) 0 % inhibition as the infection values of two consecutive high dilutions not inhibiting virus entry. We used 15 rat heart samples that represented the range of ELISA IgG ODs that we observed; we used 10 rat lung samples that included 7 ELISA IgA putatively positive samples, and 3 putatively negative samples. The tissue fluid samples were prepared as described above for the serological analysis, except that they were prepared in phosphate buffered saline only and inactivated at 56 °C for 30 mins to destroy complement or residual virus. Samples were assayed in a doubling dilution series starting from a 1 in 8 and 1 in 16 dilution for the lung and heart samples, respectively. Rat positive controls were heart and lung tissue fluid prepared from positive control SARS-CoV-2 Spike protein immunised rats (above); rat negative controls were heart and lung tissue fluid prepared from unimmunised laboratory rats. The human positive controls was serum for an individual who had had multiple SARS-CoV-2 vaccinations; the human negative control was serum from the same individual before SARS-CoV-2 vaccination.

## Results

### Animal trapping

We trapped a total of 45 rats (*Rattus norvegicus*), 10 (7 male, 3 female) from an urban park and 35 (14 male, 16 female, 5 not determined) from the sewage treatment works, and 69 mice (*A. sylvaticus*) (36 male, 33 female) from urban parks in Liverpool between 15^th^ June 2020 and 20^th^ November 2020. Our period of trapping coincided with periods of high and low SARS-CoV-2 prevalence in the people in Liverpool and northwest England more generally (**Figure 1**). This varying rate of human infection across our trapping period should therefore temporally vary the SARS-CoV-2 exposure risk to the rodents that we sampled.

### Metagenomic viral discovery

We used two SARS-CoV-2 positive controls, a high (10^8^) and a low (10^4^) dose of viral particles per gram of tissue and we used the low dose positive control to set the cut-off threshold for other viral reads. The 10^4^ difference in the viral dose between the two controls is partially reflected in the sequence reads that were recovered; specifically, the proportionate number of reads from the high and low dose controls differed by a magnitude of 3.7 and 7.2 × 10^3^ for the rat and mouse controls, respectively. This therefore generally supports the use of the proportionate number of reads in a library as a semi-quantitative, comparative measure of viral load in the wild animal samples.

Once host reads had been removed there were a total of 1.4 × 10^10^ and 6.2 × 10^9^ metagenomic sequence reads for the rat and mouse samples, respectively, with this difference possibly due to the use of a bacterial rRNA depletion bait in the mouse sample, but not the rat samples. Among these reads, 1.7 and 3.5 % of the reads from rats and mice, respectively, were Kraken2-classified as having a viral origin.

We detected a total of 297 different viruses based on Kraken2 assignment of reads. Of these 297, 264 were only found in rats, 13 only in mice, and 20 found in both species (**Supplementary Table 1**). There were an average of 37.9 (95 % CI, 31.4-45.3) different viruses in individual rats and 4.3 in individual mice (95 % CI, 3.7-4.9), which differed significantly (*df* = 68, *z* = 25.31, p<0.001). For rats we analysed the distribution of different viruses among gut, lung and faeces from analysis of tissue specific pools of 10 rats. We found that 97 viruses were only found in lung tissue, 7 only in gut tissue and 4 only in faecal material (**Supplementary Table 2**), and therefore fewer than half of the viruses we detected had a tissue-specific association.

We counted the total number of infections in rats and mice caused by different viruses and then grouped these into taxonomic families, thereby showing the viral families causing the largest proportion of infections (**Table 2**). This shows quite different representation of viral families causing infections in the two host species, with only four viral families (Picornaviridae, Baculoviridae, Poxviridae, Astroviridae) within the top ten named viral families that are shared among mice and rats. We did not detect any viruses belonging to the Coronaviridae in mice, but 0.7 % of viral infections in rats were from the Coronaviridae. It is notable that our sample processing (*i*.*e*. preparation of RNA and sequencing) has recovered reads from both RNA and DNA viruses. Viruses belonging to the Virgaviridae and Baculoviridae are present in rats, but these families are viruses of plants and arthropods, and so their presence in rats is likely to be from rodent food and / or contamination during the rodent trapping and then sample processing.

**Table 2.** The percentage of all infections caused by viral species belonging to different viral families, for the 10 highest ranked named viral families, and Coronoaviridae, showing the viral type and known host groups.

| Rat |  |  | Mouse |  |  |
| --- | --- | --- | --- | --- | --- |
| Family | Notes | Percent | Family | Notes | Percent |
| Picornaviridae | RNA; vertebrates | 16 | Peribunyaviridae | RNA; animals | 42 |
| Unclassified | - | 12 | Baculoviridae | DNA; arthropods | 20 |
| Virgaviridae | RNA; plants | 9 | Poxviridae | DNA; animals | 8 |
| Herpesviridae | DNA; vertebrates | 7 | Astroviridae | DNA; vertebrates | 7 |
| Baculoviridae | DNA; arthropods | 6 | Picornaviridae | RNA; vertebrates | 5 |
| Poxviridae | DNA; animals | 6 | Picobirnaviridae | RNA; mammals | 5 |
| Nodaviridae | RNA; animals | 6 | Hantaviridae | RNA; mammals | 4 |
| Retroviridae | RNA; animals | 4 | Unclassified | - | 3 |
| Dicistroviridae | RNA; invertebrates | 4 | Phenuiviridae | RNA; mammals | 2 |
| Astroviridae | DNA; vertebrates | 4 | Adenoviridae | DNA; vertebrates | 1 |
| Flaviviridae | RNA; vertebrates | 4 | Anelloviridae | DNA; vertebrates | 1 |
| Coronaviridae | RNA; vertebrates | 0.7 | Coronaviridae | RNA; vertebrates | 0 |

We calculated the prevalence of different viruses, considering only host samples that were not pools of different individuals. In mice three viruses (Shamonda orthobunyavirus, Simbu orthobunyavirus [both family Peribunyaviridae], Choristoneura fumiferana granulovirus [family Baculoviridae]) had a greater than 50 % prevalence, whereas in rats there were 23 viruses with a greater than 50 % prevalence, with 4 each belonging to the Virgaviridae and Picornaviridae families, and 3 to the Nodaviridae family (**Supplementary Table 3**).

Considering viral load, measured as the proportion of total reads assigned to individual viruses, for rats and mice separately, in rats 4 viruses each accounted for more than 0.5 % of reads: Rosavirus B, Boolarra virus, Norway rat hunnivirus, Cardiovirus C (**Supplementary Table 4**). In mice three viruses each accounted for more than 1 % of reads: Shamonda orthobunyavirus, Choristoneura fumiferana granulovirus, Simbu orthobunyavirus. For rats and mice the viruses with these large number of reads were also among the viruses that were most prevalent (above).

### Coronaviridae and SARS-CoV-2

Three rats had SARS-CoV-2 sequence reads whose proportionate number was greater than the low dose positive control (which was 10^4^ viral particles per gram of tissue). The three positives were pools of material; specifically, 1 pool of lung tissue from 3 rats, 2 pools of gut tissue each from 3 different rats; these three pools encompass 7 rats caught in an urban park in Liverpool in July 2020. When reads from these samples were aligned by Bowtie2 to the SARS-CoV-2 genome, 8, 39 and 72 read pairs aligned from the three samples (compared to 72 read pairs for low dose positive control) and their starting positions typically corresponded with primer sites from the ARTIC SARS-CoV-2 primer scheme used in the laboratory as part of the UK COVID surveillance programme, and so these reads likely represent low level contamination. In contrast, and as expected, reads from the positive controls had a uniform pattern of alignment across the genome. No reads aligned to the SARS-CoV-1 genome. Together, these metagenomic data indicate that we did not detect natural SARS-CoV-2 infection in rats. No mouse samples had SARS-CoV2 reads greater than the relevant positive controls.

### Other Coronaviridae

We detected coronavirus-identified metagenomic sequence reads (above our threshold) in rats; specifically Ferret coronavirus (an alphacoronavirus) in 7 rat samples (4 individuals and 3 pools of lung tissue each from 3 or 4 rats); Rhinolophus bat coronavirus HKU2 (an alphacoronavirus) in one rat sample (a pool of lung tissue from 3 rats), and Middle East respiratory syndrome-related coronavirus (a betacoronavirus) in 7 rat samples (6 individuals and 1 pool of gut tissue from 3 rats). We were unable to assemble genome sequence of these three viruses.

We did not detect any coronaviruses metagenomic sequence reads (above our cut-off threshold) in mice. However, we did notice that many mouse samples had substantial numbers of sequence reads identified as coming from an Avian coronavirus (a gammacoronavirus). No single mouse sample had more reads than our threshold cut-off, though by halving the threshold, then 9 mouse samples were positive (5 individual samples and 2 pools of pairs of mice).

### Hantaviridae

We found evidence of hantaviruses within the rat and mouse samples. Specifically. Oxbow orthohantavirus in 16 rat samples (10 individuals, 3 pools of lung tissue and 3 pools of gut tissue each from 3 or 4 rats) and in 6 mouse samples (all individuals), and Seoul orthohantavirus in one rat individual sample. For the Seoul orthohantavirus we were able to partially assemble the sequence, with the largest contig of 4.8 kb and an N50 of 1.5 kb; BLAST analysis of this confirmed the identity as Seoul orthohantavirus.

### SARS-CoV-2 – PCR detection

We did not PCR detect SARS-CoV-2 in the wild rat and mouse samples. Our high positive control strongly amplified, our low positive control amplified less strongly, and our negative controls did not amplify. Therefore, these data provide evidence of the absence of SARS-CoV-2 RNA in our wild rat and mouse samples.

### SARS-CoV-2 – Serology

We first tested our ability to detect immunoglobulin in tissue fluid samples. Specifically, we found that negative control laboratory rats’ heart and liver tissue fluid had a concentration of 166 and 437 μg / mL of total IgG, respectively, compared with a concentration of 6,770 μg / mL in serum. Heart and liver samples therefore represent 2.4 and 6.4 % of the IgG concentration in serum. A randomly selected wild rat had similar concentrations of total IgG in heart and liver tissue fluid, specifically 670 and 400 μg / mL, respectively. These concentrations of total IgG in a wild rat are therefore the same order of magnitude as those found in laboratory animals, though higher concentrations may be expected in wild animal due to their higher antigenic exposure (Abolins *et al*., 2017). Concerning antibodies to the SARS-CoV-2 Spike protein, in positive control, Spike protein immunised animals the titre was 640,000 in serum and 32,000 in liver and heart tissue fluid samples. These results for the relative concentration of antigenic specific antibodies in serum *vs*. tissue are compatible with those for the concentration of total IgG. Together these results show that it is possible to detect IgG antibodies and antigen-specific antibodies in tissue fluid samples, as well as in serum, although the observed concentration is lower in tissue fluid samples compared with serum.

For total IgA, in positive control, Spike protein immunised rats we detected IgA in serum at a concentration of 301 μg / mL and at lower concentrations in tissue fluid, specifically 28, 7.9 and 1.5 μg / mL in lung, heart and liver, respectively. In two wild rats there were high concentrations of total IgA in lung tissue fluid, 307 and 1,188 μg / mL. We suspect that the comparatively very high concentration of IgA in wild rat lungs is because of their high antigenic exposure in the wild (Abolins *et al*., 2017). These results show that it is possible to detect IgA in lung tissue fluid.

Based on these results we then screened rat and mouse heart tissue fluid samples for anti-SARS-CoV-2 Spike IgG antibodies and rat lung tissue fluid for anti-SARS-CoV-2 Spike IgA antibodies.

In rats we used a 1:640 dilution of heart tissue fluid in which we detected very low concentrations of anti-SARS-CoV-2 Spike IgG antibodies, with the OD of the wild rats being less than a third of the positive control (**Figure 2**). We used a 1:20 dilution of lung tissue fluid to assay for anti-SARS-CoV-2 Spike IgA antibodies, similarly finding that most animals had very low concentrations, though 7 rats had ODs greater than or equal to the positive control (**Figure 3**). In interpreting this result it is important to remember that the positive control animals were immunised intramuscularly, and not in the lung and in this respect our positive control presumably does not maximise the lung IgA response. Notwithstanding, the OD of 7 wild rats are high, and there are two interpretations of these data. First, that this is evidence of a lung IgA response produced by rats following exposure to and / or infection with SARS-CoV-2. The second, alternative explanation is that these lung IgA antibodies are cross-reactive to SARS-CoV-2 Spike protein, but produced in response to an infection with a different virus. Of these 7 rats, 6 are samples from southern England donated by pest controllers; the remaining rat was caught in an urban park in Liverpool in January 2020.

**Figure 2.**
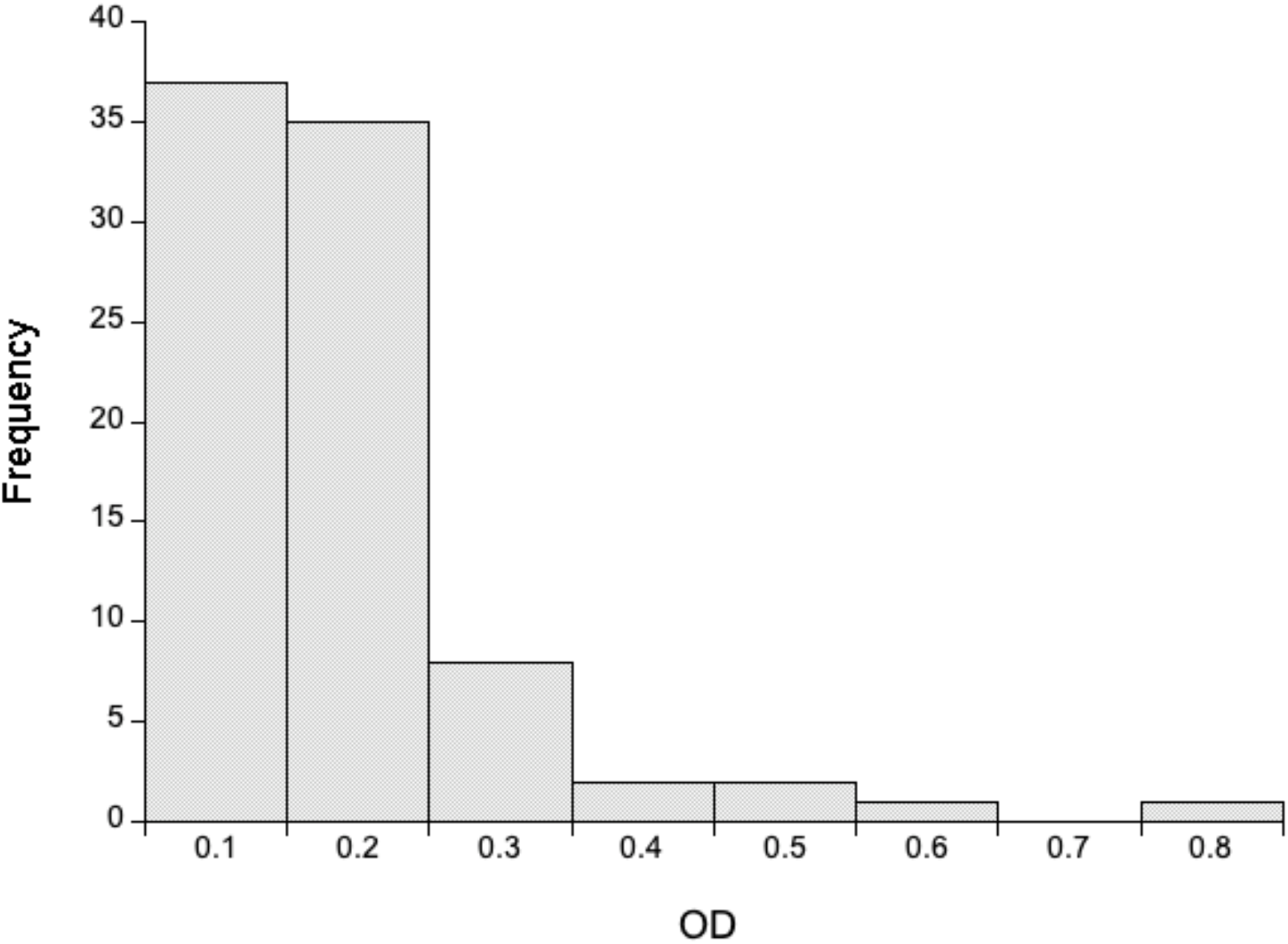
The distribution of OD of anti-SARS-CoV-2 IgG responses for 86 wild rats for 1:640 diluted heart tissue fluid. The OD of the positive control heart tissue fluid is 3.55.

**Figure 3.**
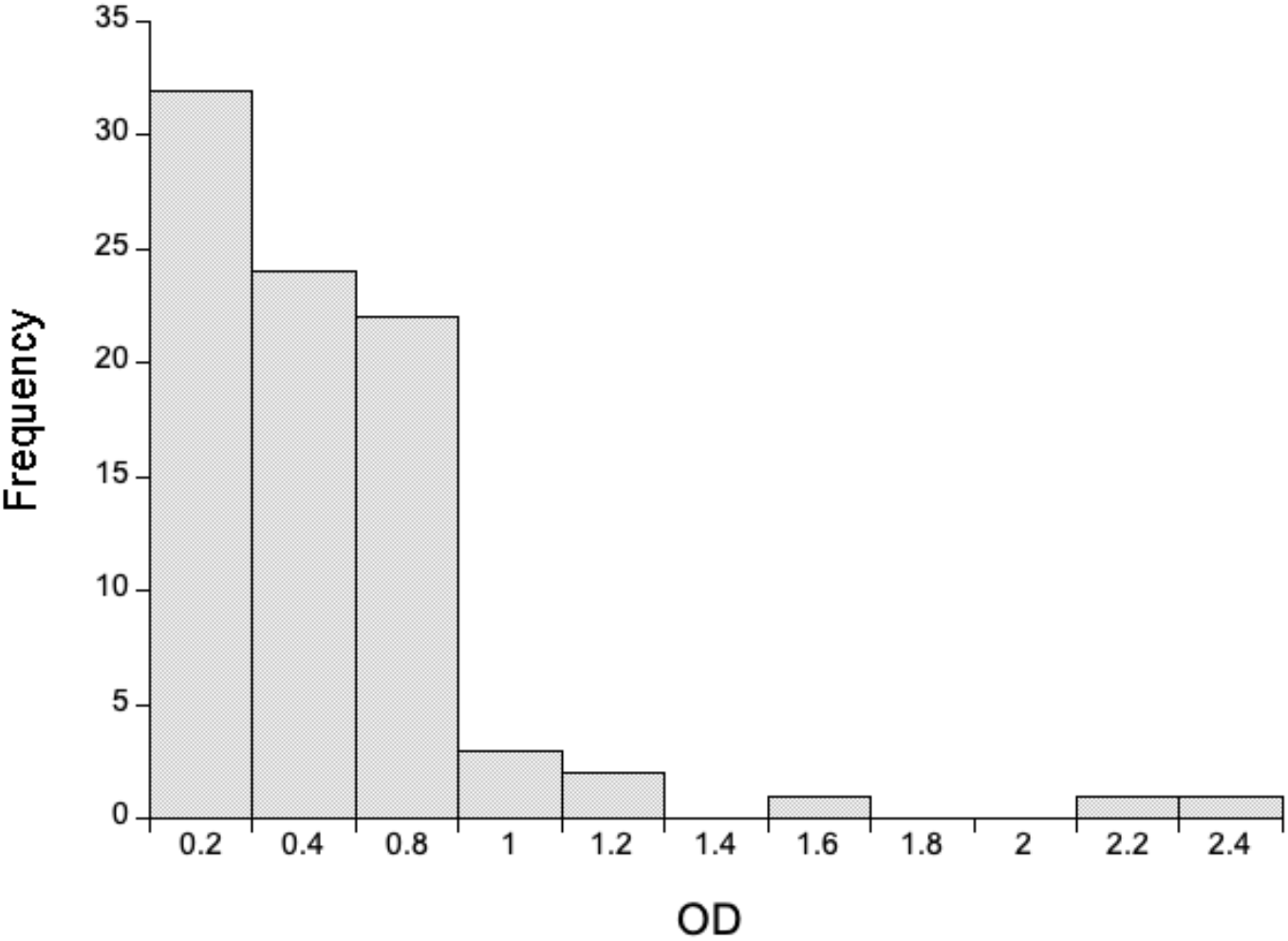
The distribution of OD of anti-SARS-CoV-2 IgA responses for 86 wild rats for 1:20 diluted lung tissue fluid. The OD of the positive control lung tissue fluid is 0.825.

In mice we used a 1:160 dilution of heart tissue fluid in which we detected very low concentrations of anti-SARS-CoV-2 Spike IgG antibodies, with the OD of the wild rats generally a tenth of the positive control (**Figure 4**). In interpreting these data it is important to remember that the positive control is a *M. musculus* samples, while the wild mouse is *A. sylvaticus*, but that the ELISA used *M. musculus* reagents.

**Figure 4.**
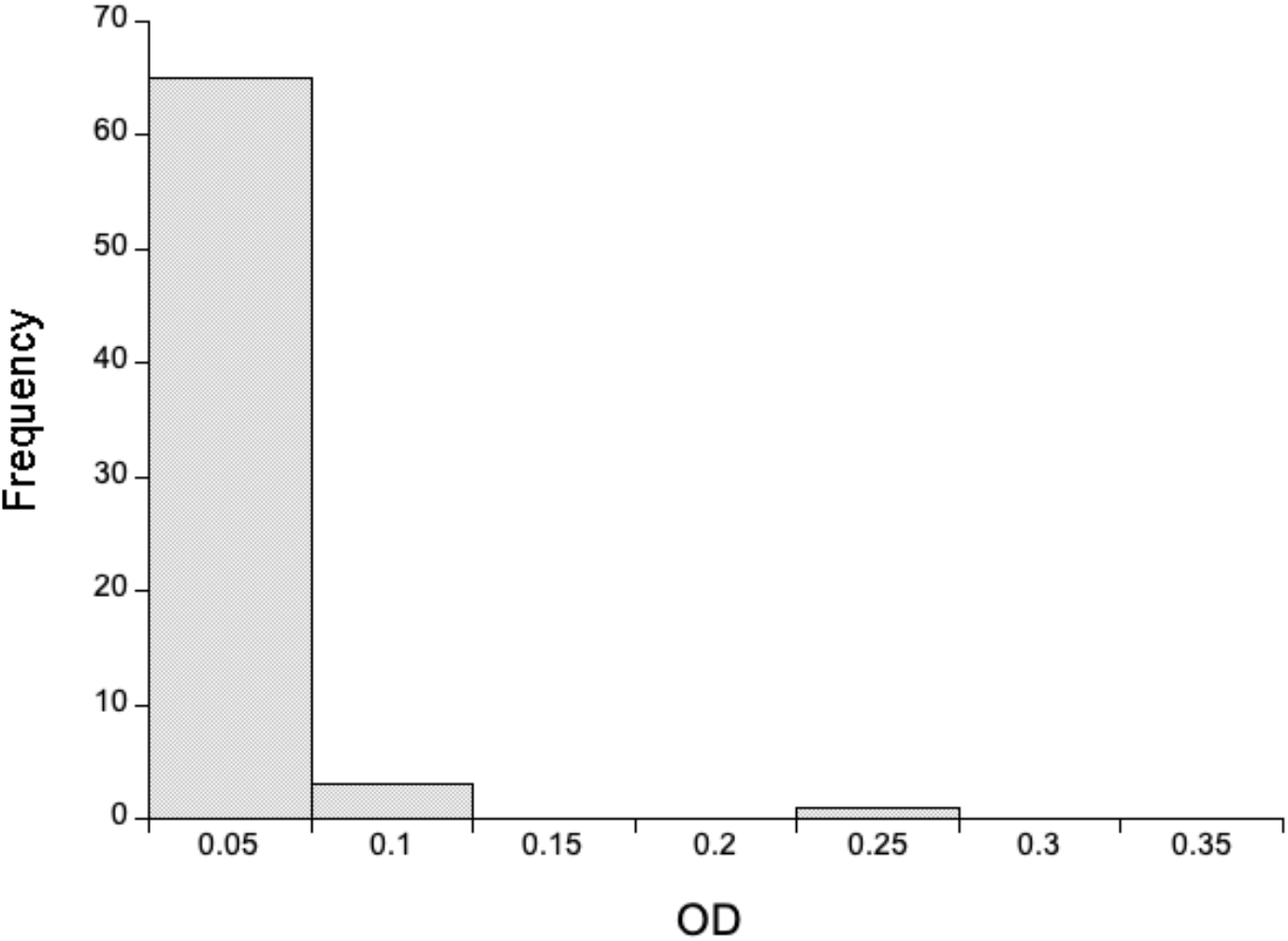
The distribution of OD of anti-SARS-CoV-2 IgG responses for 69 wild mice for 1:160 diluted heart tissue fluid. The OD of the positive control heart tissue fluid is 0.532.

### Neutralization assays

We tested the ability of rat heart and lung tissue fluid to neutralize SARS-CoV-2 pseudovirus particles’ ability to infect HEK293T ACE2 TMPRSS2 cells. We found that three (of 15) rat heart samples partially neutralized pseudovirus particle infection (36-70 % inhibition) at the highest concentration of tissue fluid, compared with 84 % neutralization by the rat positive control (**Figure 5**). There was no obvious relationship between the ELISA IgG ODs and the neutralization. For the rat lung samples, four samples achieved 39 – 61 % neutralization at the highest concentration of tissue fluid, compared with 93% neutralization by the positive control (**Figure 5**). Three of these four samples were putatively IgA positive samples. Of particular note, lung and heart tissue from a single rat achieved 59 and 23 % neutralization, respectively. These results are consistent with the idea that wild rats are being exposed to SARS-CoV-2 and generating an immune response that has the ability to neutralise infection *in vitro*. The alternative interpretation is that wild rats are exposed to coronaviruses other than SARS-CoV-2 and that antibodies to these other viruses are partially neutralising. These interpretations are not mutually exclusive.

**Figure 5.**
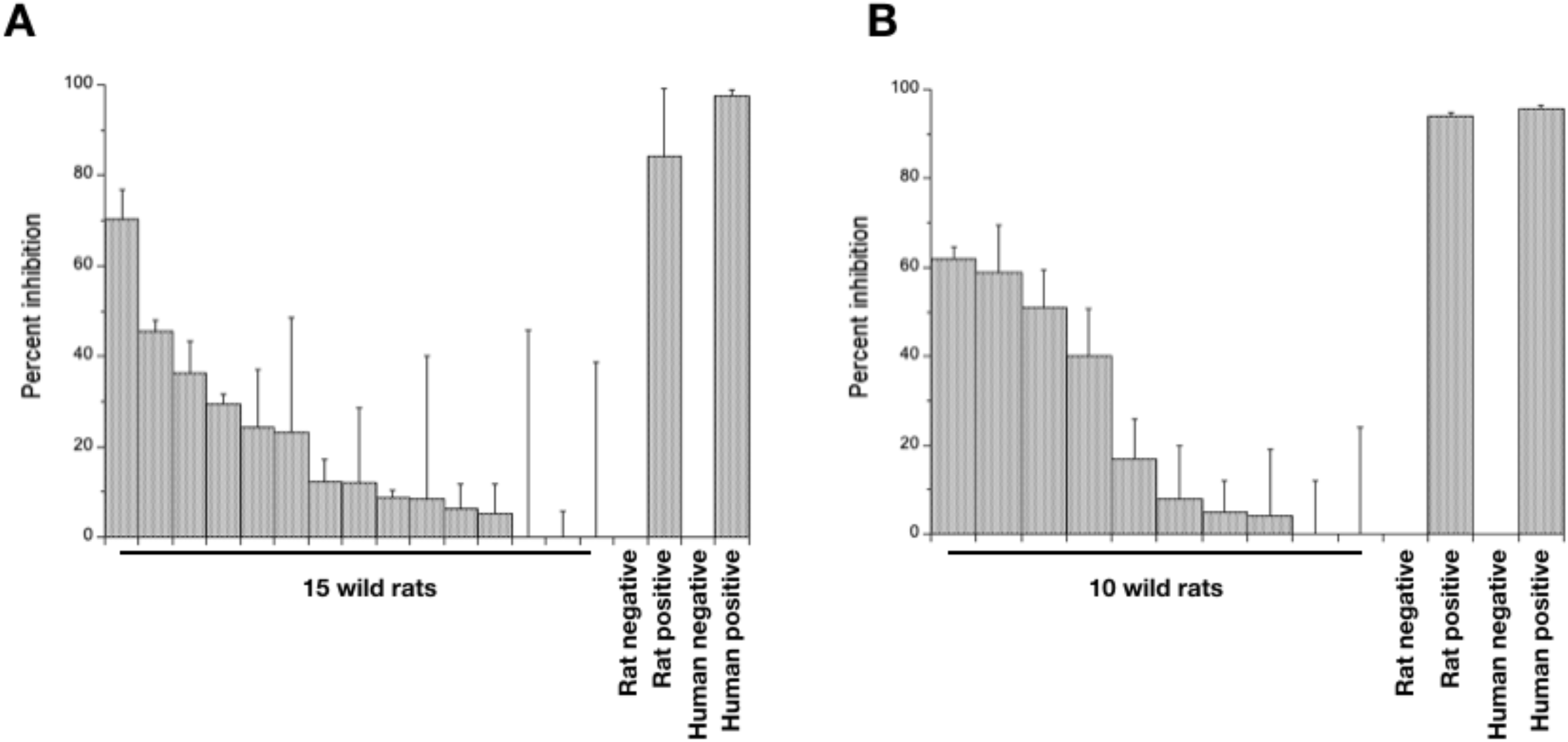
The mean percent inhibition of wild rat (A) heart and (B) lung tissue fluid at a 1 in 16 and 1 in 8 dilution, respectively. Errors bars are +1 SD. Rat and human positive and negative controls are as described in the Materials and Methods.

## Discussion

We have investigated the viruses present in urban rat and mouse populations, with a specific focus on determining whether or not they are exposed to and / or infected with SARS-CoV-2. In investigating this we were testing the hypothesis of a reverse zoonosis of SARS-CoV-2 from infected people to rodents during the 2020 human pandemic. In this we were mindful of a range of different scenarios, ranging from full, long-lived infections in rodents, through short-term infections, to exposure that didn’t result in a patent infection. Using a combination of PCR-based detection of SARS-CoV-2, metagenomic sequencing of lung, gut and faeces, and serological analysis we find some evidence consistent with rats being exposed to SARS-CoV-2. We found no such evidence for mice.

For rats, the evidence is (i) lung anti-SARS-CoV-2 Spike IgA responses equal to or greater than our positive control and (ii) that lung and heart tissue fluid is able to partially neutralise SARS-CoV-2 PVP *in vitro*. Our interpretation of these data is that rats are being exposed to SARS-CoV-2, perhaps resulting in short-lived infections that results in detectable anti-SARS-CoV-2 antibodies. The alternative interpretation of these data is (i) that the IgA antibodies were generated in response to other viral infections and that these antibodies cross-react with SARS-CoV-2 Spike protein, and (ii) that these and other antibodies generated in response to other viral infections can partially neutralise SARS-CoV-2.

Further studies will be needed to separate these possibilities. Our results are therefore consistent with studies of rats in Belgium and Hong King where there was evidence of antibodies that recognised SARS-CoV-2 and with some evidence of their ability to neutralise the virus (Colombo *et al*., 2021; Miot *et al*., 2022). Our approach goes further than these studies by considering lung IgA responses, which may be more appropriate in understanding rodent exposure to SARS-CoV-2 compared with serum IgG.

In a scenario where rats are occasionally exposed to low doses of SARS-CoV-2, then it would be challenging to detect this molecularly since most of the time rats would not have SARS-CoV-2 virus present. Moreover the presence of antibodies that recognise and neutralise SARS-CoV-2 in wild rat populations would further abrogate any potential infection, making molecular detection of the virus harder still. In our metagenomic sequencing we used positive controls of low dose SARS-CoV-2, so providing an objective evidence base for detecting the presence of true virus-derived nucleic acid.

For occasional short-lived, low dose SARS-CoV-2 infection, serological analyses are potentially very useful, because any immunological response to such infection or exposure will persist through the life of an animal. The challenge with interpreting serological data are two-fold. First, cross-reaction of antibodies that were produced in response to other, non-SARS-CoV-2, viruses thus producing false positive results. Second, the absence of appropriate positive controls for use in serological assays. Our serology positive controls were laboratory animals immunised with purified SARS-CoV-2 Spike protein in adjuvant, twice, intramuscularly. This results in a high anti-SARS-CoV-2 antibody titre, but such a titre is unlikely to ever be achieved in a natural infection of a wild animal. Therefore a better positive control would be a pulmonary or enteric infection of low doses of virus, which would then be appropriate to the type and magnitude of infection that we hypothesise may be occurring in wild rodents. It is also important to remember that the immune state of wild animals is quite different to that of laboratory animals (Abolins *et al*., 2017), further limiting the utility of laboratory animal infections in producing positive controls appropriate for use in studying wild animals.

A UK DEFRA risk assessment of SARS-CoV-2 infection of rodents concluded with satisfactory evidence that there was a high likelihood that SARS-CoV-2 VOC could infect a commensal rodent, and the data we provide here is potential evidence of this. This risk assessment went on to consider knowledge gaps, one of which was the viral dose required to infect a rodent and the degree of viral shedding from any such infected animal, but our results do not provide evidence to address this.

We also used our metagenomic sequence analysis to identify other viruses in rats and mice. In this, our low dose positive control provided an objective cut-off for determining a positive signal of viral infection. Comparison of the high and low dose controls also provided evidence that the number of viral reads can be used as a semi-quantitative measure of viral abundance. We found evidence of almost 300 viruses, principally in rats, and then mainly from lung tissue. We used a Kraken2 database to identify these viruses, which is good initial evidence of viral identity, though assembly of viral genome sequence is better evidence of viral identity. In this respect our identification of viruses should be considered indicative rather than definitive. We identified sequence reads from a wide range of viral families, though the most highly represented families differed between rats and mice. These results are broadly comparable to other molecular surveys of viruses in wild rodents, including: in the US a range of viruses already known from mammals (including members of the Coronaviridae, Astroviridae, Picornaviridae, Picobirnaviridae, Adenoviridae, Papillomaviridae, Parvovirinae and Circoviridae) as well as viruses known from insects and plants; in gut samples from rats in Berlin, Germany a wide range of viruses (Phan *et al*., 2011; Sachsenröder *et al*., 2014). Other molecular surveys of viruses in rodents (*Apodemus* spp. and *Myodes glareolus*) has found that much of the faecal virome is seasonally transient, emphasising the need for longitudinal studies (Raghwani *et al*., 2022). However, overall together these studies show that wild rodents commonly have a rich and diverse virome.

We also found evidence of other coronavirus infection in our samples; specifically, in rats of a betacoronavirus, Middle East respiratory syndrome-related coronavirus (MERS) and two alphacoronaviruses; in mice a gammcoronavirus. Coronaviruses have previously been detected in wild rodents, including: alphacoronaviruses in UK rats (but not in *Mus* spp. or *Apodemus* spp.) (Tsoleridis *et al*., 2016; Tsoleridis *et al*., 2019); betacoronaviruses in *Apodemus* sp. in France (Monchatre-leroy *et al*., 2017); alphacoronaviruses in rodents in the Congo basin (Kumakamba *et al*., 2021); alpha and betacoronaviruses in *Apodemus* spp. and *Rattus* spp. in China (Wang *et al*., 2015; Wang *et al*., 2020); coronaviruses in house mice (*M. musculus*) but not in *Rattus* spp. in the Canary Islands (Monastiri *et al*., 2021), and a high seroprevalence against coronaviruses in US *R. norvegicus* (Easterbrook *et al*., 2007). Together, this shows that a range of coronaviruses, including betacoronaviruses, do occur in wild rodents, which contributes to addressing the DEFRA-identified knowledge gaps of the endogenous coronaviruses of rodents. The potential for recombination of SARS-CoV-2 with such endogenous coronaviruses (also a DEFRA-identified knowledge gap) remains unknown, but the data we present is supportive of rats being potentially exposed to SARS-CoV-2 that are infected with other coronaviruses is the necessary prelude to any potential recombination.

Concerning our putative identification of MERS in rats, MERS is related to a number of bat coronaviruses and to a hedgehog coronavirus (Corman *et al*., 2014). The hedgehog coronavirus is relatively common in hedgehogs in Europe with a prevalence ranging from 10 – 58 % (Saldanha, *et al*., 2019; Delogu *et al*., 2020; Pomorska-Mól *et al*., 2022). The Kraken2 database we used to identify our metagenomic sequence reads contains both MERS, hedgehog coronavirus, and other bat coronaviruses. On balance we suspect that our rat MERS viral reads are more likely to be derived from a hedgehog coronavirus, or perhaps a bat coronavirus, rather than *sensu stricto* MERS since (i) both hedgehogs and bats live within the environments where we caught rats and (ii) that MERS has only been reported to infect humans, bats and camels (Widagdo *et al*., 2019) The other alternative is that these rats MERS-assigned reads are derived from a hitherto unknown betacoronavirus. To resolve this, definitive identification of the rat virus would be needed, which would require viral genome assembly, which we were unable to achieve. Notwithstanding, the discovery of an additional betacoronavirus in rats leads credence to the idea that there could be recombination among betacoronavirsues in rats. These result address further knowledge gaps identified by the DEFRA risk assessment, specifically (i) the endogenous coronaviruses of rodents and (ii) the risk of recombination of SARS-CoV-2 with other coronaviruses. In mice we found good evidence for the presence of Avian gammacoronavirus.

We found evidence of hantavirus infection in rats and mice, specifically of Oxbow othohantavirus in rats and mice and Seoul orthohantavirus in a rat. Hantaviruses are common in rodent hosts, and if humans become infected this can cause two syndromes: haemorrhagic fever with renal syndrome and cardiopulmonary syndrome (Vaheri *et al*., 2013). Seoul virus has a worldwide distribution and in the UK has been found in wild and pet rats (Dupiney *et al*., 2014; McElhinney *et al*., 2017; Ling *et al*., 2019; Murphy *et al*., 2019), and so our report is consistent with these previous descriptions. Oxbow orthohantavirus was first described from an American shrew mole (*Neurotrichus gibbsii*) (King *et al*., 1976), after which there are no further reports. We have found evidence of the widespread presence of this virus in rats and mice (*A. sylvaticus*), and if this is substantiated then this would be a significant extension of the know host and geographical range of this virus.

Our metagenomic analysis identified quite different viromes in rats and mice, notably with the rat virome being much larger (284 viruses in rats *vs*. 33 in mice). Our processing of animals, RNA preparation and sequencing of material from these two host species was done at different times and the rRNA removal included bacterial baits in the mouse samples, but not in the rat samples. In all this resulted in comparatively more reads in the rat samples compared with the mouse samples (1.4 × 10^10^ *vs*. 6.2 × 10^9^, respectively), but with a two -fold greater proportion of these being identified as viral reads in mice than in rats (3.5 *vs*. 1.7 %, respectively). Therefore there seems to be a contradiction between the high rat *vs*. low mouse viral diversity we report compared with the high-level view of the sequence data. We used rat and mouse positive controls to set objectively a cut-off threshold. We suspect that this cut-off may have been too high in the mouse samples; if this is correct then lowering of this threshold would result in us reporting many more viruses in mice.

In conclusion, we present evidence consistent with rats being exposed to SARS-CoV-2, possibly from a human source. If substantiated, this could have important implications for the future evolutionary trajectory and epidemiology of SARS-CoV-2 and for the future risk of viral recombination with potential risk for animal and human populations.

### Data Deposition

The metagenomic sequence data are deposited in the European Nucleotide Archive as accessions PRJEB53828 and PRJEB5329.

## Authors’ contributions

**AF** planned and conducted the field work, contributed to the laboratory work, data analysis and interpretation, and co-wrote the paper;

**GA** planned and conducted the laboratory work, contributed to data analysis and interpretation, and co-wrote the paper;

**YL** contributed to the laboratory work, to data analysis and interpretation, and co-wrote the paper;

**MG** undertook the bioinformatic analysis and interpretation, and co-wrote the paper;

**EB** contributed to the laboratory work, to data analysis and interpretation, and co-wrote the paper;

**MW** contributed to protocol design, data analysis and interpretation, and co-write the paper;

**JT** conducted the neutralization assays, contributed to data analysis and interpretation, and co-wrote the paper;

**WP** co-conceived the study, contributed to study design and data interpretation, and co-wrote the paper;

**GP** co-conceived the study, contributed to study design and data interpretation, and co-wrote the paper;

**SP** co-conceived the study, contributed to study design and data analysis and interpretation, and co-wrote the paper;

**MV** conceived and led the study, contributed to data analysis and interpretation, and co-wrote the paper;

## Acknowledgments

We would like to thank Mark Garth, Adam Johnson and colleagues for assistance with trapping; Alex Wade for help in obtaining rodent samples; David Hall, Simon King and Andy Brigham from the pest control industry for help and advice; Sam Haldenby for bioinformatics advice; Eleanor Riley for advice and discussions; James Stewart for help with the control immunisations. This work was funded by a grant from NERC and funds from the University of Liverpool. EGB is supported by the University of Liverpool; YL is supported by the University of Liverpool and the China Scholarship Council.

## Supplementary Tables

**Supplementary Table 1.**
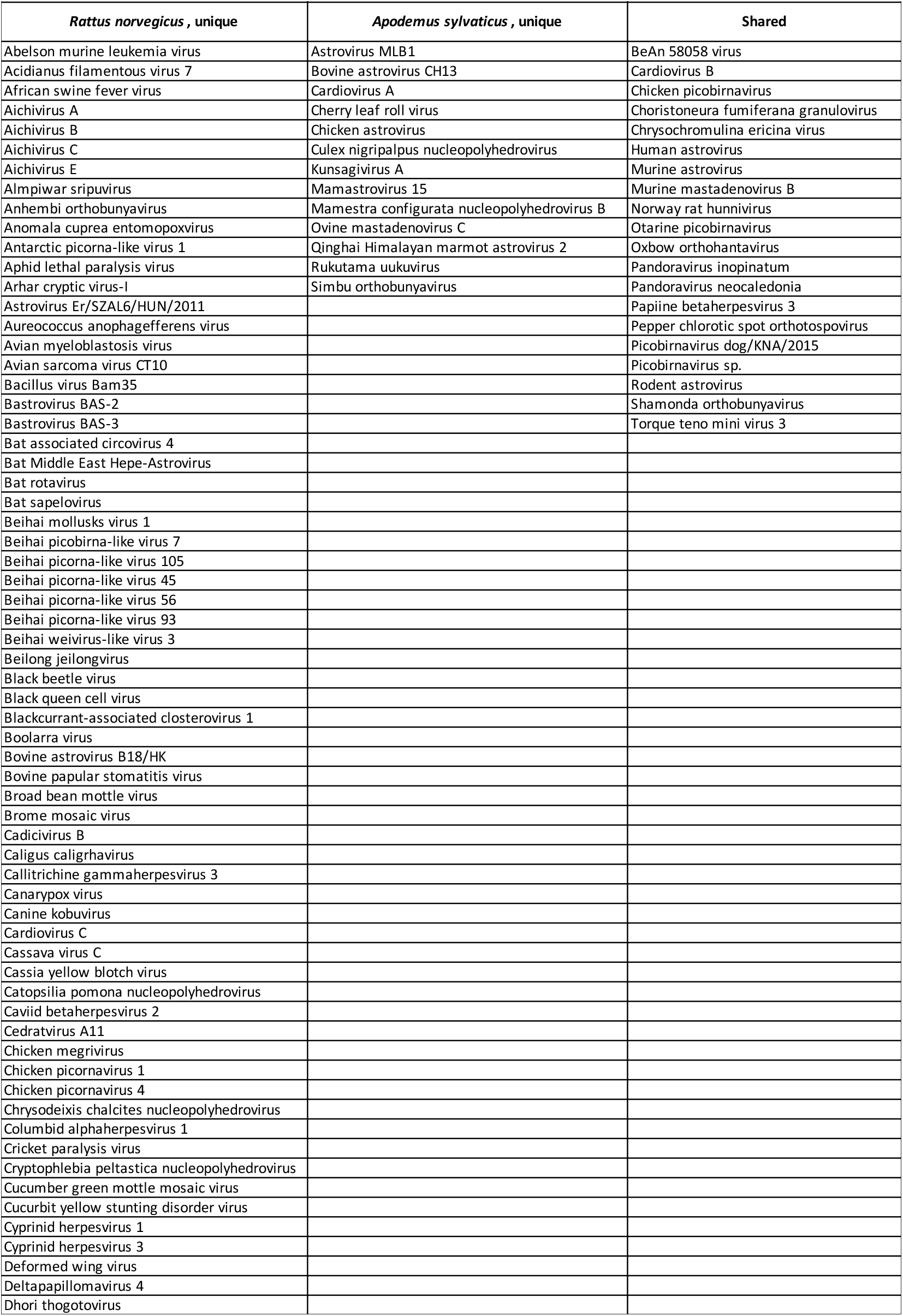
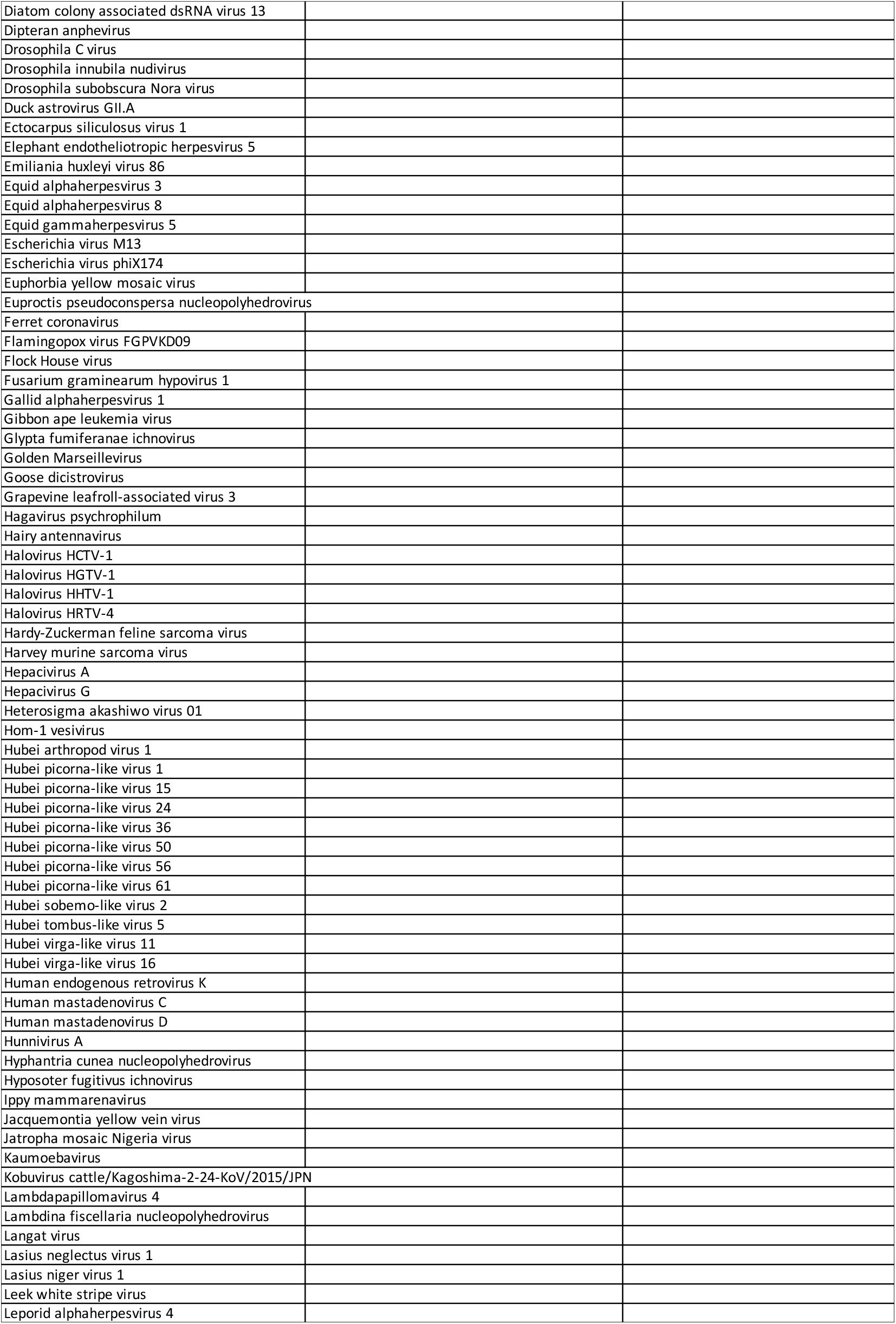
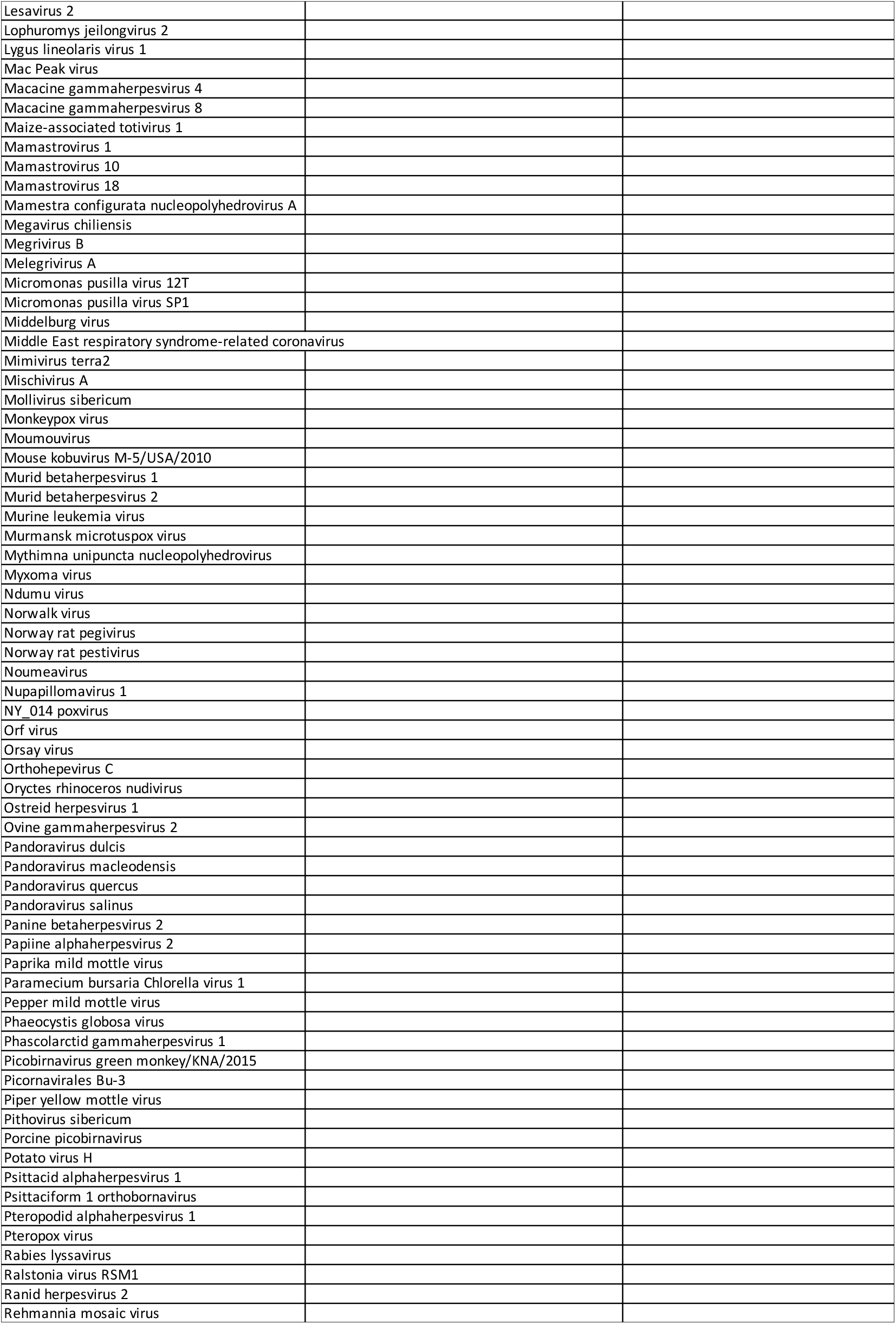
Viruses unique to *Rattus norvegicus* or *Apodemus sylvaticus*, or shared between then, based on Kraken2-assigend sequence reads.

**Supplementary Table 2.**
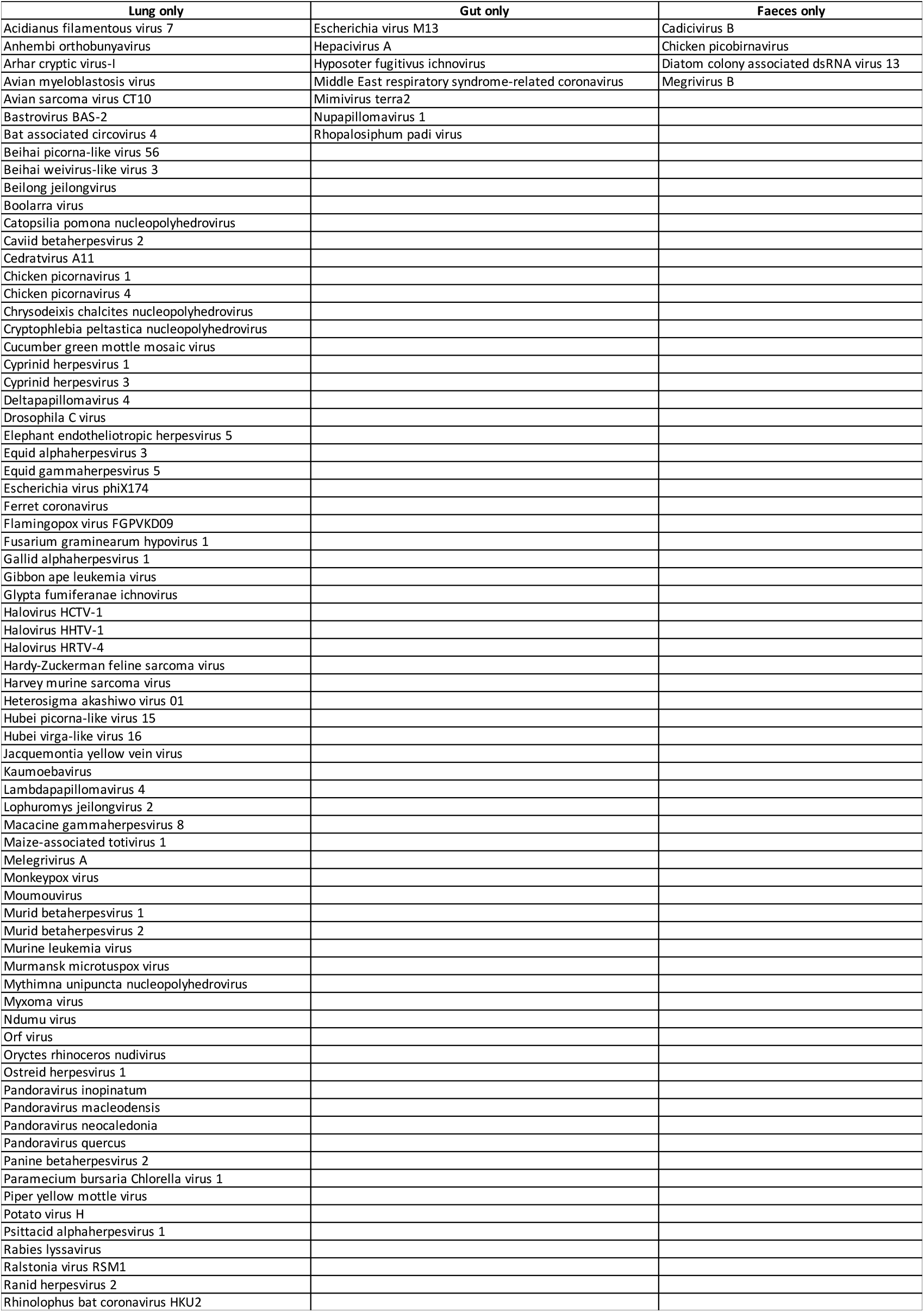
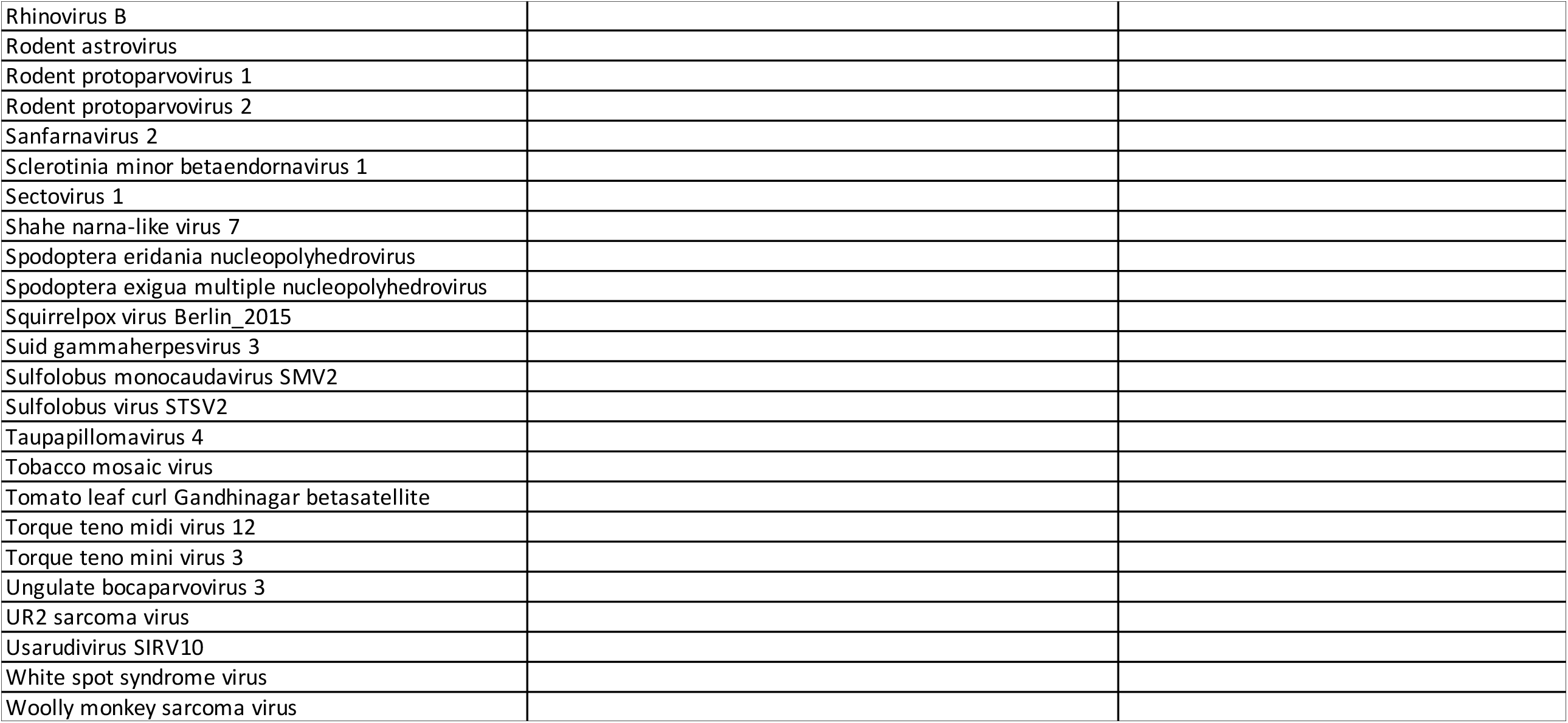
The viruses found only in lung tissue, gut tissue of faecal material of rats.

**Supplementary Table 3.**
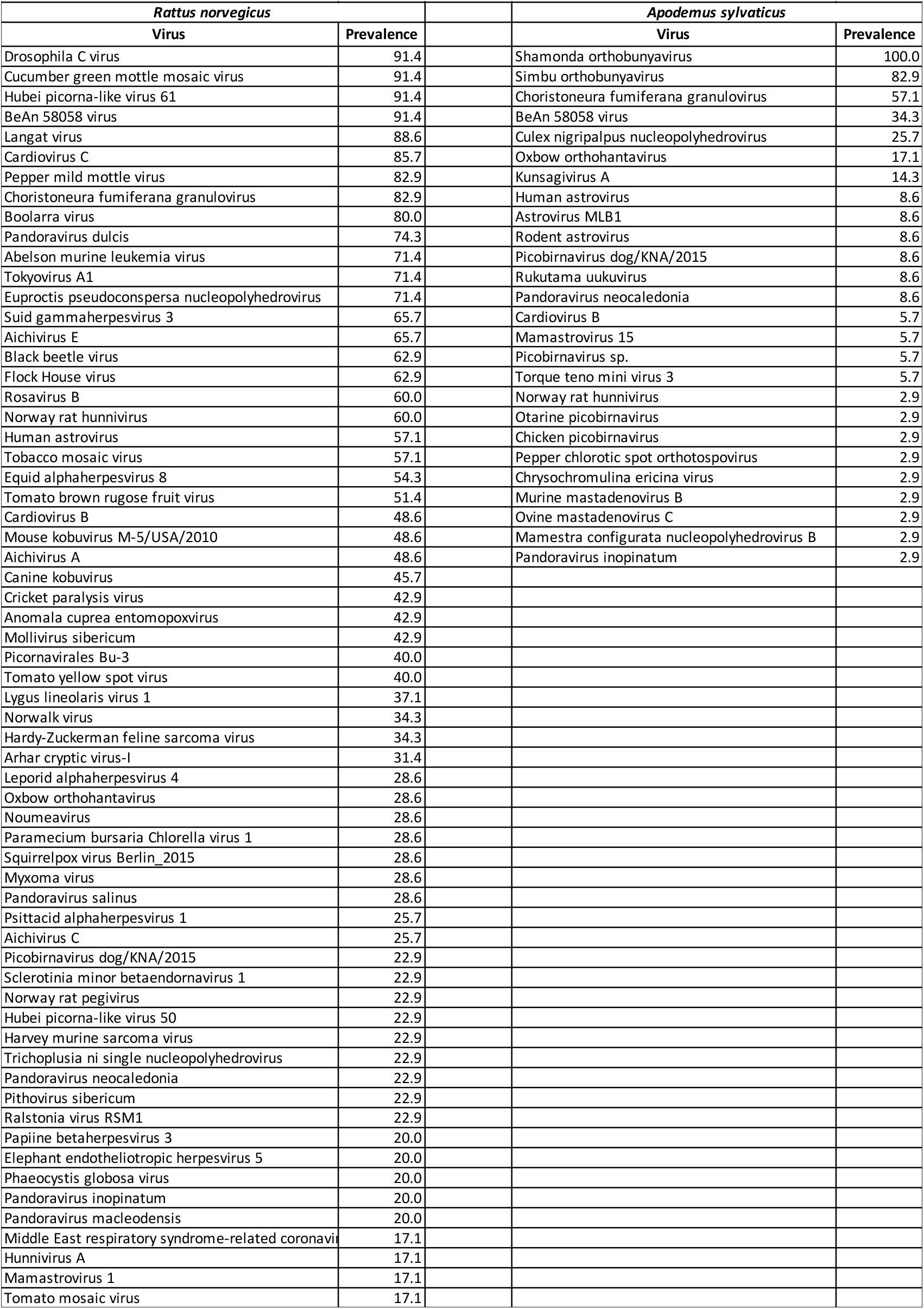
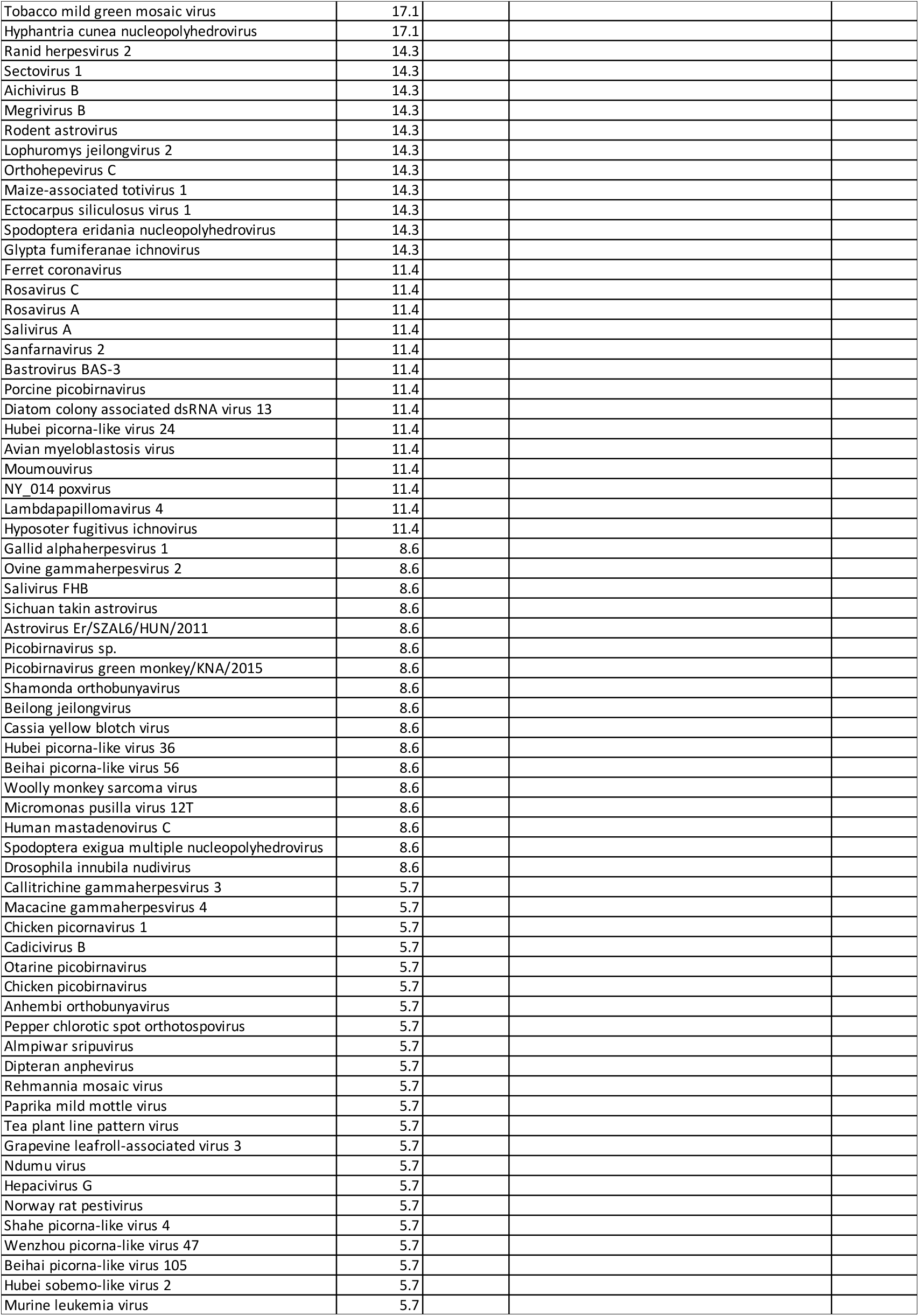
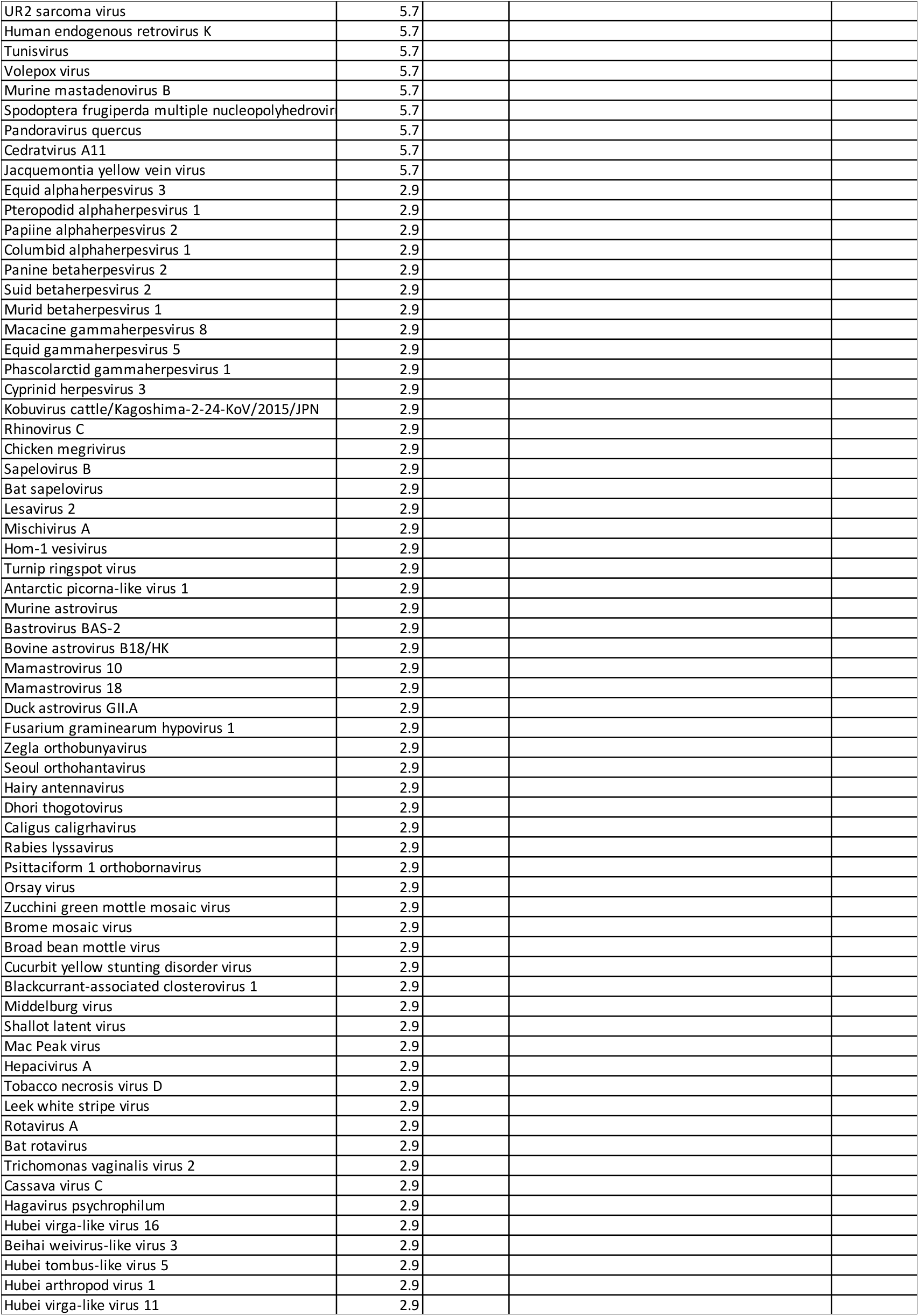
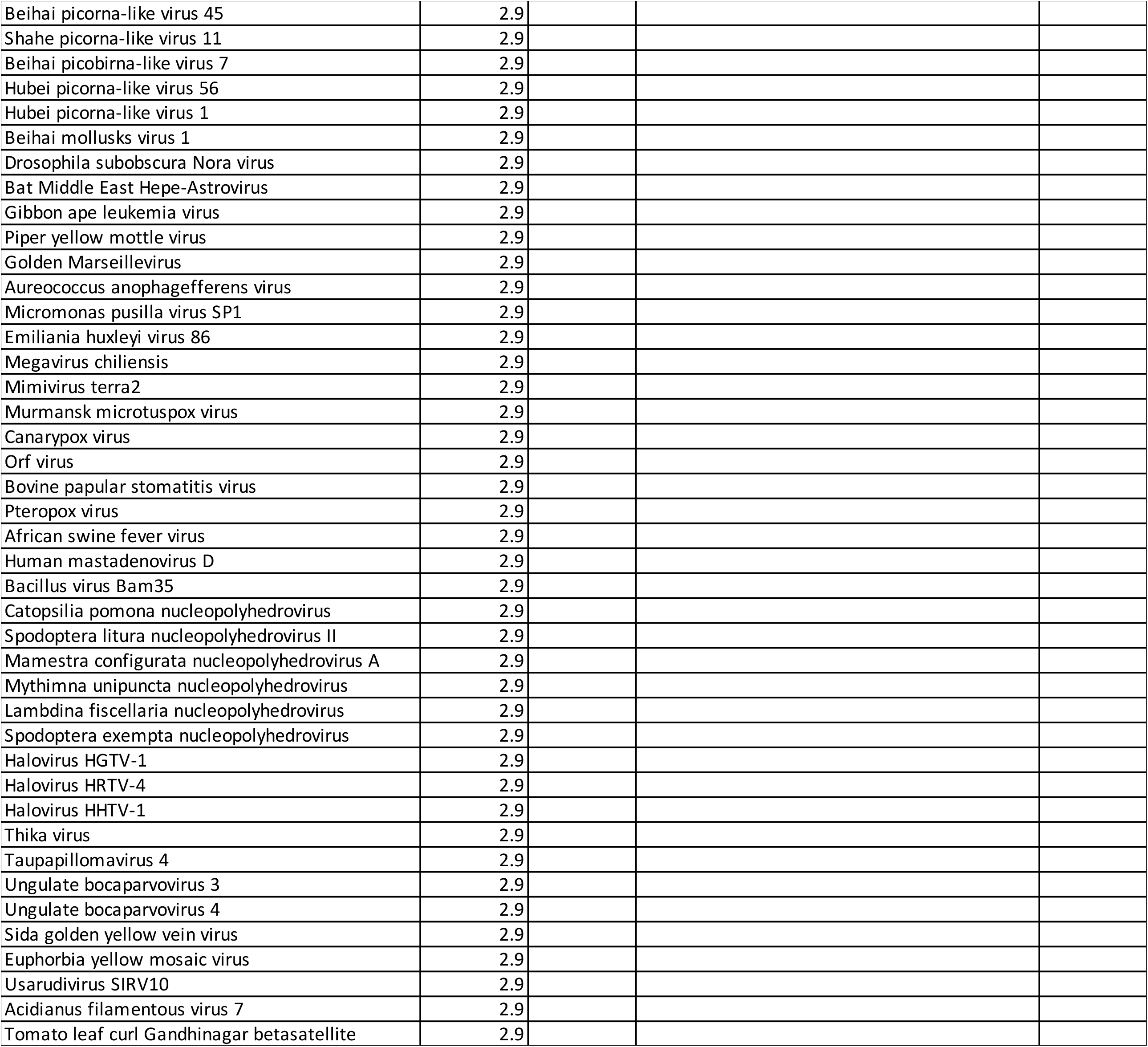
The prevalence of different viruses in *Rattus norvegicus* and *Apodemus sylvaticus*.

**Supplementary Table 4.**
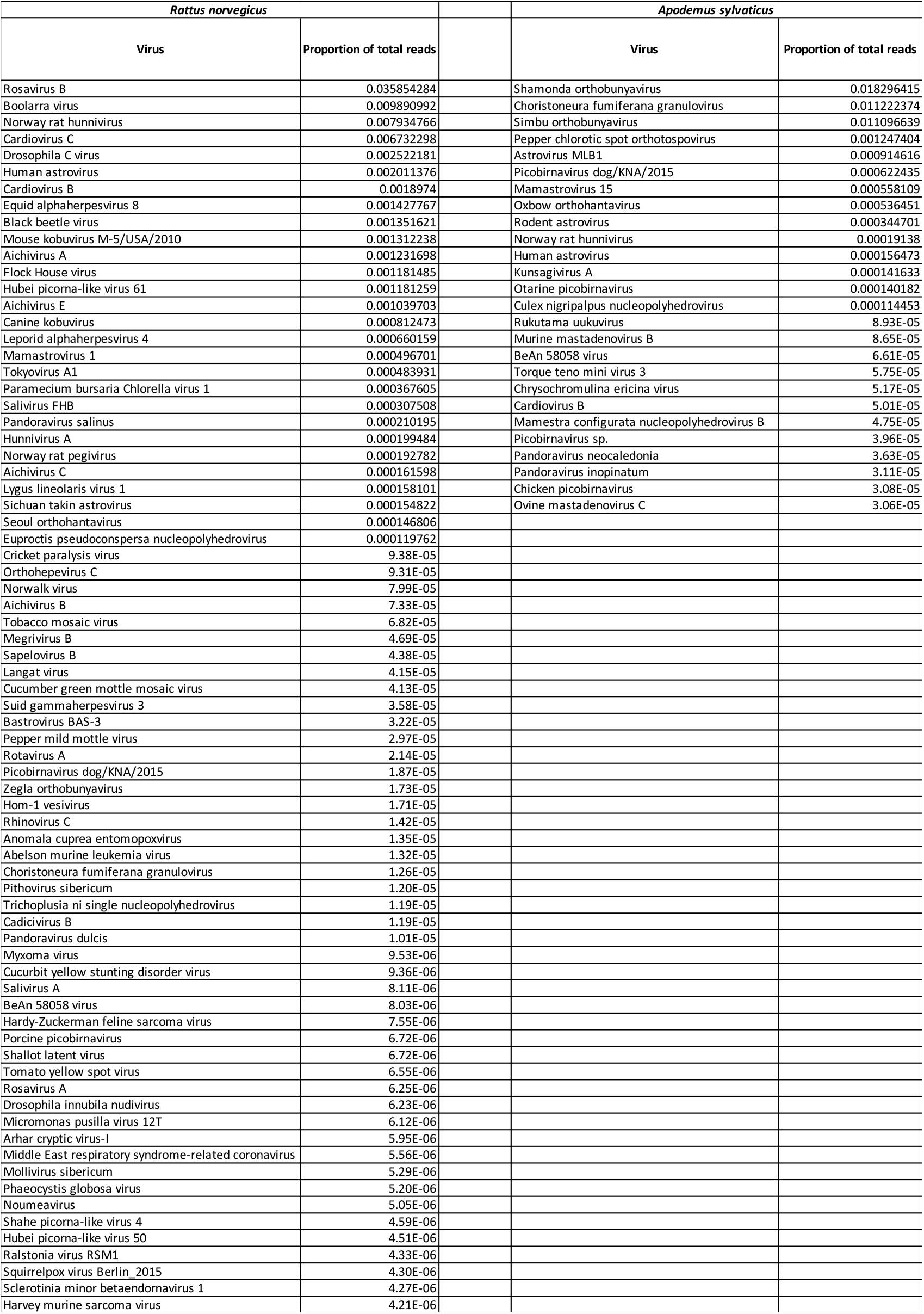
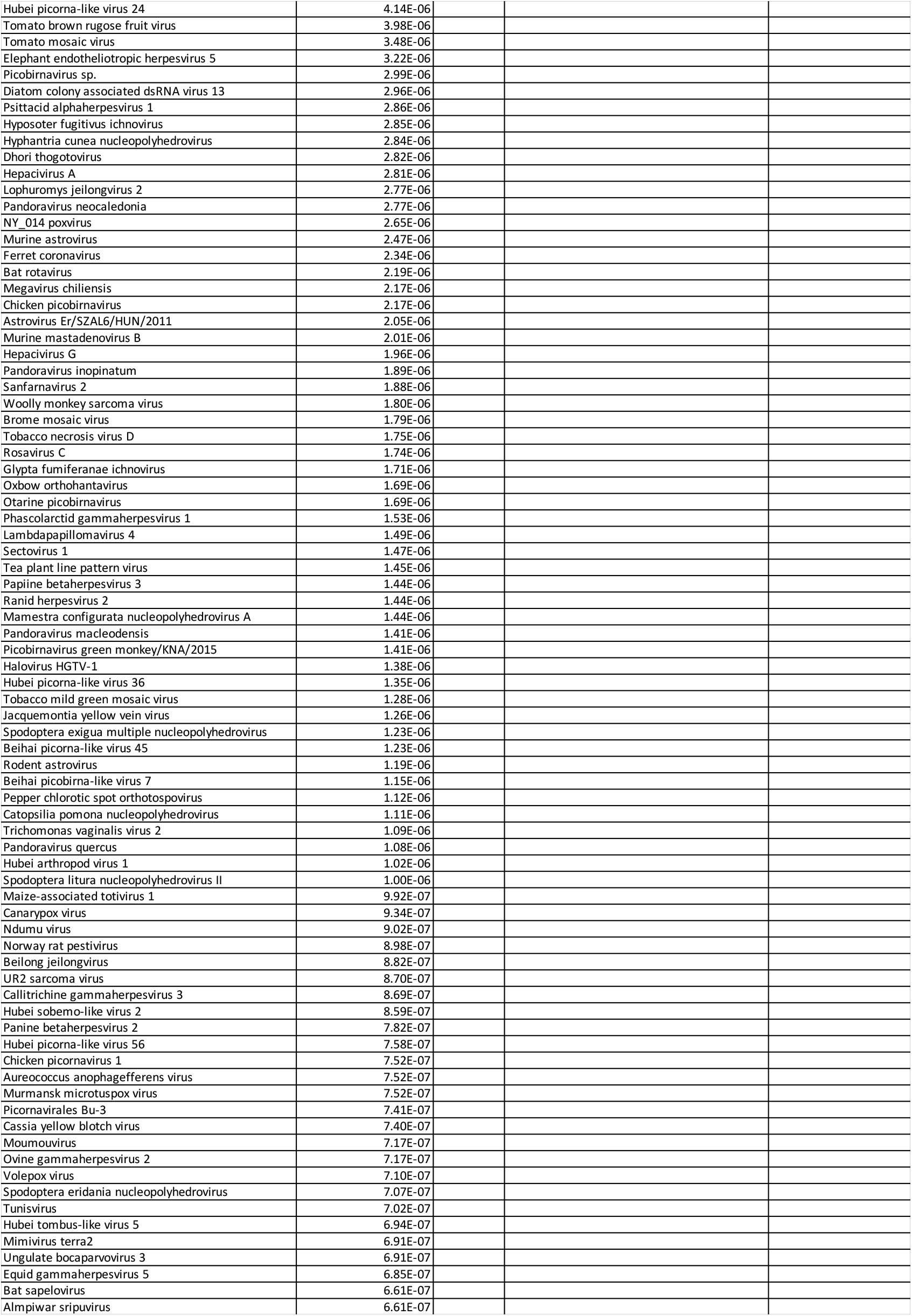
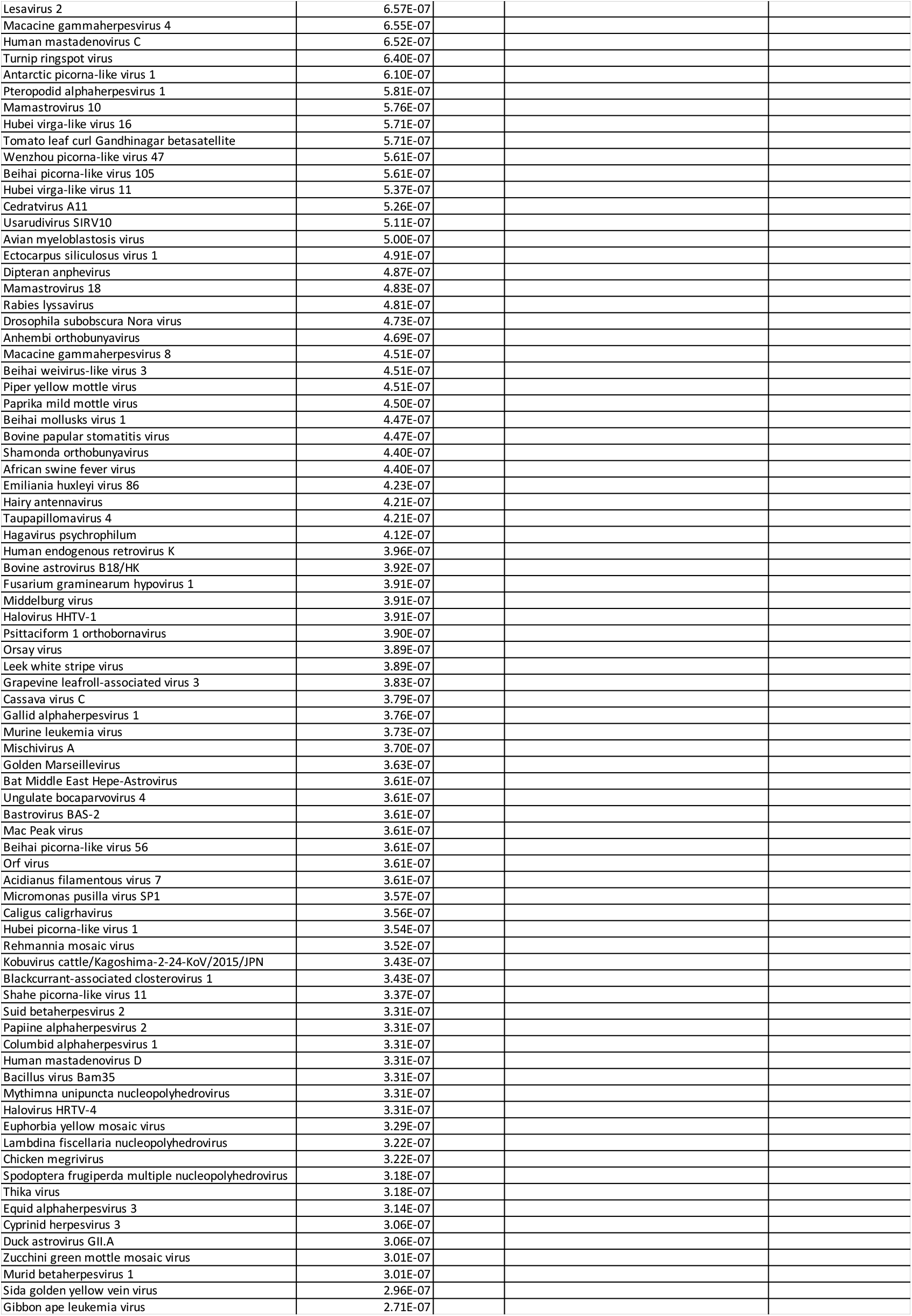
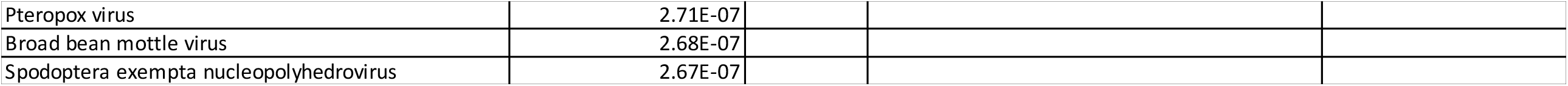
The viral load, measured as the proportion of total reads assigned to each virus, in *Rattus norvegicus* and *Apodemus sylvaticus*.

## References

Abolins, S., King, E., Lazarou, L., Weldon, L., Hughes, L., Drescher P., et al., 2017. The comparative immunology of wild and laboratory mice Mus musculus domesticus. Nat. Commun. 8, 14811.

Adaken, C., Scott, J.T., Sharma, R., Gopal, R., Dicks, S., Niazi, S., et al. (2021). Ebola virus antibody decay–stimulation in a high proportion of survivors. Nature 590, 468–472.

Albery, G.F., Carlson, C.J., Cohen, L.E., Eskew, E.A., Gibb, R., Ryan, S.J. et al., 2022. Urban-adapted mammal species have more known pathogens. Nat. Ecol. Evol. 6, 794–801.

Artic 2020. https://github.com/artic-network/primer-schemes/blob/master/nCoV-2019/V3/nCoV-2019.tsv

Bankevich, A., Nurk, S., Antipov, D., Gurevich, A.A., Dvorkin, M., Kulikov, et al. 2012. SPAdes: a new genome assembly algorithm and its applications to single-cell sequencing. J. Comp. Biol. 19, 455–477.

Brown, M.R., Wade, M.J., McIntrye-Nolan, S., Bassano, I., Denise, H., Bass, D et al., 2020. Wastewater monitoring of SARS-CoV-2 variants in England: Demonstration case study for Bristol (Dec 2020 – March 2021). Summary for SAGE 8th April 2021. https://www.gov.uk/government/publications/jbc-and-defra-wastewater-monitoring-of-sars-cov-2-variants-in-england-demonstration-case-study-for-bristol-december-2020-to-march-2021-8-april-20

Capizzi, D., Bertolino, S., Mortelliti, A., 2014. Rating the rat: global patterns and research priorities in impacts and management of rodent pests. Mammal Rev. 44, 148–162.

Carnell, G., Grehan, K., Ferrara, F., Molesti, E., Temperton, N. 2017. An optimized method for the production using PEI, titration and neutralizationof SARS-CoV Spike Luciferase Pseudotypes. Bio Protoc. 7, e2514.

Chan, J.F., Zhang, A.J., Yuan, S., Poon, V.K., Chan, C.C., Lee, A.C., et al., 2020. Simulation of the clinical and pathological manifestations of coronavirus disease 2019 (COVID-19) in a golden syrian hamster model: Implications for disease pathogenesis and transmissibility. Clin. Infect. Dis. 71, 2428–2446.

Chen, L., Liu, B., Wu, Z., Jin, Q., Yang, J., 2017. DRodVir: A resource for exploring the virome diversity in rodents. J. Genet. Genomics 44, 259–264.

Clerc, M., Babayan, S.A., Fenton, A., Pedersen, A.B., 2019. Age affects antibody levels and anthelmintic treatment efficacy in a wild rodent. Int. J. Parasitol. Parasites Wildl. 8, 240–247.

Colombo, V.C., Sluydts, V., Mariën, J., Vanden Broecke, B., Van Houtte, N., Leirs, W. et al., 2021. SARS-CoV-2 surveillance in Norway rats (Rattus norvegicus) from Antwerp sewer system, Belgium. Transbound Emerg Dis. 2021, 1–6. (doi: 10.1111/tbed.14219)

Corman, V. M., Kallies, R., Philipps, H., Göpner, G., Müller, M.A., Eckerle. I, et al., 2014. Characterization of a novel betacoronavirus related to middle East respiratory syndrome coronavirus in European hedgehogs. J. Virol. 88, 717–724

DEFRA 2021. https://www.gov.uk/government/publications/jbc-and-defra-a-qualitative-risk-assessment-to-estimate-the-likelihood-of-sars-cov-2-infection-of-rodents-from-contact-with-the-environment-and-onwar

Delogu, M., Cotti, C., Lelli, D., Sozzi, E., Trogu, T., Lavazza, A., et al., 2020. EcovVirological preliminary study of potentially emerging pathogens in Hedgehogs (Erinaceus europaeus) recovered at a wildlife treatment and rehabilitation center in northern Italy. Animals 10, 407.

Di Genova, C., Sampson, A., Scott, S., Cantoni, D., Mayora-Neto, M., Bentley, E., et al. 2021 Titration, neutralisation, storage and lyophilisation of Severe Acute Respiratory Syndrome Coronavirus 2 (SARS-CoV-2) Lentiviral Pseudotypes. Bio Protoc., 11, e4236.

Dupinay, T., Pounder, K.C., Ayral, F., Laaberki, M.H., Marston, D.A., Lacôte, S., et al., 2014. Detection and genetic characterization of Seoul virus from commensal brown rats in France. Virol. J. 11, 32.

Easterbrook, J.D., Kaplan, J.B., Glass, G.E., Watson, J., Klein, S.L. 2008. A survey of rodent-borne pathogens carried by wild-caught Norway rats: A potential threat to laboratory rodent colonies. Lab Animal 42, 92–98.

Fagre, A., Lewis, J., Eckley, M., Zhan, S., Rocha, S.M., Sexton, N.R., et al., 2021. SARS-CoV-2 infection, neuropathogenesis and transmission among deer mice: Implications for spillback to New World rodents. PLoS Pathog. 17, e1009585.

Ge, X.Y., Yang, W.H., Zhou, J.H., Li, B., Zhang, W., Shi, Z-L., et al., 2017. Detection of alpha-and betacoronaviruses in rodents from Yunnan, China. Virol. J. 14, 98.

Gibb, R., Redding, D.W., Chin, K.Q., Donnelly, C.A., Blackburn, T.M., Newbold, T., Jones, K.E., et al., 2020. Zoonotic host diversity increases in human-dominated ecosystems. Nature 584, 398–402.

Gu, H., Chen, Q., Yang, G., He, L., Fan, H., Deng, Y.Q., et al., 2020. Adaptation of SARS-CoV-2 in BALB/c mice for testing vaccine efficacy. Science 369, 1603–1607.

Han, B.A., Schmidt, J.P., Bowden, S.E., Drake, J.M., 2015. Rodent reservoirs of future zoonotic diseases. Proc. Natl. Acad. Sci. USA 112, 7039–7044.

Hassell, J.M., Begon, M., Ward, M.J., Fèvre, E.M., 2017. Urbanization and disease emergence: dynamics at the wildlife–livestock–human interface. Trends Ecol. Evol. 32, 55–67.

Huang, K., Zhang, Y., Hui, X., Zhao, Y., Gong, W., Wang, T., et al., 2021. Q493K and Q498H substitutions in Spike promote adaptation of SARS-CoV-2 in mice. EbioMed. 67, 103381.

Huang, H., Zhu, Y., Niu, Z., Zhou, L., Sun, Q. 2021. SARS-CoV-2 N501Y variants of concern and their potential transmission by mouse. Cell Death Differ. 28, 2840–2842.

Jackson, J.A., Friberg, I.M., Bolch, L., Lowe, A., Ralli, C., Harris, P.D., et al., 2009. Immunomodulatory parasites and toll-like receptor-mediated tumour necrosis factor alpha responsiveness in wild mammals. BMC Biol. 7, 16.

Janik, E., Niemcewicz, M., Podogrocki, M., Majsterek, I., Bijak, M., 2021. The emerging concern and interest SARS-CoV-2 variants. Pathogens 10, 633.

Karthikeyan, S., Levy, J.I., De Hoff, P., Humphrey G., Birmingham A., Jepsen, K., et al., 2022. Wastewater sequencing reveals early cryptic SARS-CoV-2 variant transmission. Nature 609, 101–108.

Kumakamba, C., Niama, F.R., Muyembe, F., Mombouli, J.V., Kingebeni, P.M., Nina, R.A. et al., 2021. Coronavirus surveillance in wildlife from two Congo basin countries detects RNA of multiple species circulating in bats and rodents. PLoS One 16m e0236971.

Langmead B, Salzberg S. 2012. Fast gapped-read alignment with Bowtie 2. Nat. Meth. 9, 357–359.

Lau, S.K., Woo, P.C., Li, K.S., Tsang, A.K., Fan, R.Y., Luk, H.K., et al., 2015. Discovery of a novel coronavirus, China Rattus coronavirus HKU24, from Norway rats supports the murine origin of Betacoronavirus 1 and has implications for the ancestor of Betacoronavirus lineage A. J. Virol. 89, 3076–3092.

Ling, J., Verner-Carlsson, J., Eriksson, P., Plyusnina, A., Löhmus, M., Järhult, J.D., et al., 2019. Genetic analyses of Seoul hantavirus genome recovered from rats (Rattus norvegicus) in the Netherlands unveils diverse routes of spread into Europe. J. Med. Virol. 91, 724–730.

Martin, M., 2011. Cutadapt removes adapter sequences from high-throughput sequencing reads. EMBnet J. 17, 1.

McElhinney, L.M., Marston, D.A., Pounder, K.C., Goharriz, H., Wise, E.L., Verner-Carlsson, J., et al., 2017. High prevalence of Seoul hantavirus in a breeding colony of pet rats. Epidemiol. Infect. 145, 3115–3124.

Miot, E.F., Worthington, B.M., Ng, K.H., de Lataillade, L.D.G., Pierce, M.P., Liao, Y., et al., 2022. Surveillance of rodent pests for SARS-CoV-2 and other Coronaviruses, Hong Kong. Emerg. Infect. Dis. 28, 467–470.

Monastiri, A., Martín-Carrillo, N., Foronda, P., Izquierdo-Rodríguez, E., Feliu, C., López-Roig, M., et al,. 2021. First coronavirus active survey in rodents from the Canary Islands. Front. Vet. Sci. 8, 708079.

Monchatre-Leroy, E., Boué, F., Boucher, J.M., Renault, C., Moutou, F., Ar Gouilh, M., et al., 2017. Identification of alpha and beta coronavirus in wildlife species in France: bats, rodents, rabbits, and hedgehogs. Viruses, 9, 364.

Montagutelli, X., Prot, M., Levillayer, L., Salazar, E.B., Jouvion, G., Conquet, L., et al., 2021. The B1.351 and P.1 variants extend SARS-CoV-2 host range to mice. bioRxiv 2021.03.18.436013; doi: https://doi.org/10.1101/2021.03.18.436013

Murphy, E.G., Williams, N.J., Bennett, M., Jennings, D., Chantrey, J., McElhinney, L.M. 2019. Detection of Seoul virus in wild brown rats (Rattus norvegicus) from pig farms in Northern England. Vet. Record 184, 525–525.

Otto, S.P., Day, T., Arnio, J., Colin, C., Dushoff, J., Li, M., et al., 2021. The origins and potential future of SARS-CoV-2 variants of concern in the evolving COVID-19 pandemic. Curr. Biol. 31, R918–R929.

Pomorska-Mól, M., Ruszkowski, J.J., Gogulski, M. Domanska-Blicharz, K., 2022. First detection of Hedgehog coronavirus 1 in Poland. Sci. Rep. 12, 2386.

Phan, T.G., Kapusinszky, B., Wang, C., Rose, R.K., Lipton, H.L., Delwart, E.L. 2011. The fecal viral flora of wild rodents. PLoS Pathog. 7, e1002218.

Raghwani, J., Faust, C.L., François, S., Nguyen, D., Marsh, K., Raulo, A., et al., 2022. Seasonal dynamics of the wild rodent faecal virome. bioRxiv 2022.02.09.479684; doi:https://doi.org/10.1101/2022.02.09.479684

Sachsenröder, J., Braun, A., Machnowska, P., Ng, T.F.F., Deng, X., Guenther, S., et al., 2014. Metagenomic identification of novel enteric viruses in urban wild rats and genome characterization of a group A rotavirus. J. Gen. Virol. 95, 2734–2747.

Saldanha, I.F., Lawson, B., Goharriz, H., Rodriguez-Ramos Fernandez, J., John, S.K., Fooks, A.R., et al., 2019. Extension of the known distribution of a novel clade C betacoronavirus in a wildlife host. Epidemiol. Infect. 147, e169.

Shukla, N., Roelle, S.M., Suzart, V.G., Bruchez, A.M., Matreyek, K.A. 2012. Mutants of human ACE2 differentially promote SARS-CoV and SARS-CoV-2 spike mediated infection. PLoS Pathog. 17, e1009715.

Smyth, D.S., Trujillo, M., Gregory, D.A. Cheung, K., Gao, A., Graham, M., et al., Tracking cryptic SARS-CoV-2 lineages detected in NYC wastewater. Nat. Commun. 13, 635.

Su, S., Shen, J., Zhu, L., Qiu, Y., He, J-S., Tan, J-Y., et al., 2020. Involvement of digestive system in COVID-19: manifestation, pathology, management and challenges. Therap. Adv. Gastroenterol. 13, 1–12.

Taylor, L.H., Latham, S.M., Woolhouse, M.E., 2001. Risk factors for human disease emergence. Philos. Trans. R. Soc. Lond. B Biol. Sci. 356, 983–989.

Thorvaldsdóttir, H., Robinson, J.T., Mesirov, J.P., 2013. Integrative Genomics Viewer (IGV): high-performance genomics data visualization and exploration. Brief. Bioinf. 14, 178–192.

Tsoleridis, T., Onianwa, O., Horncastle, E., Dayman, E., Zhu, M., Danjittrong, T., et al., 2016. Discovery of novel alphacoronaviruses in European rodents and shrews Viruses 8, 84

Tsoleridis, T., Chappell, J.G., Onianwa, O., Marston, D.A., Fooks, A.R., Monchatre-Leroy, E., et al., 2019. Shared common ancestry of rodent alphacoronaviruses sampled globally. Viruses 11, 125.

Vaheri, A., Strandin, T., Hepojoki, J., Sironen, T., Henttonen, H., Mäkelä, S., et al., 2013. Uncovering the mysteries of hantavirus infections. Nat. Rev. Microbiol. 11, 539–550.

Wang, W., Lin, X.D., Guo, W.P., Zhou, R.H., Wang, M.R., Wang, C.Q., et al., 2015. Discovery, diversity and evolution of novel coronaviruses sampled from rodents in China. Virology 474, 19–27.

Wang, W., Lin, X.D., Zhang, H.L., Wang, M.R., Guan, X.Q., Holmes, E.C., et al., 2020. Extensive genetic diversity and host range of rodent-borne coronaviruses. Virus Evol. 6, veaa078.

Wardeh, M., Baylis, M., Blagrove, M.S., 2021. Predicting mammalian hosts in which novel coronaviruses can be generated. Nat. Commun. 12, 1–12.

Widagdo, W., Sooksawasdi Na Ayudhya, S., Hundie, G.B., Haagmans, B.L. 2019. Host determinants of MERS-CoV transmission and pathogenesis. Viruses 11, 280.

Wood, D. E., Lu, J., Langmead, B., 2019. Improved metagenomic analysis with Kraken 2. Genome Biol. 20, 257.

Woolhouse, M.E., Taylor, L.H., Haydon, D.T., 2001. Population biology of multihost pathogens. Science 292, 1109–1112.

Woolhouse, M.E., Gowtage-Sequeria, S., 2005. Host range and emerging and remerging pathogens. Emerg. Infect. Dis. 11, 1842.

Wu, Y., Guo, C., Tang, L., Hong, Z., Zhou, J., Dong, X., et al., 2020. Prolonged presence of SARS-CoV-2 viral RNA in faecal samples. Lancet Gastroenterol. Hepatol. 5, P434–P435.

Yang, J., Wang, W., Chen, Z., Lu, S., Yang, F., Bi, Z., et al., 2020. A vaccine targeting the RBD of the S protein of SARS-CoV-2 induces protective immunity. Nature 586, 572–577.

